# Human frontoparietal cortex represents behaviorally-relevant target status during invariant object recognition

**DOI:** 10.1101/387498

**Authors:** Margaret Henderson, John T. Serences

## Abstract

Searching for items that are useful given current goals, or “target” recognition, requires an observer to generalize across identity-preserving transformations such as viewpoint changes, as well as to incorporate contextual information. While past work has found target recognition signals in areas of ventral visual cortex, it is not clear whether these signals support performance on demanding tasks that require invariant, flexible search. Here, we used a task that required subjects to match novel object stimuli based on invariant features (identity and viewpoint). Based on multivariate fMRI analyses, the data suggest that the multiple-demand (MD) network, including sub-regions of parietal and frontal cortex, encodes invariant representations of an object’s status as a target. Furthermore, target information in MD regions, but not early or ventral visual cortex, was higher on correct compared to incorrect trials, suggesting a strong link between MD target signals and behavior.

## Introduction

Humans can hold information about relevant objects in mind for long periods of time, despite the presence of distracting visual inputs, and can use this stored information to flexibly guide behavior. This ability is critical for everyday search tasks, in which a viewer needs to identify a target item based on its similarity to an item held in working memory. In the simplest case, this task can be accomplished by comparing simple features of the template and viewed items (e.g., finding a photograph based on memory of the same photograph). However, this comparison problem is much more complex under natural viewing conditions, where the retinal projection of a sought object can have considerable variability due to changes in pose, position, and environmental conditions (DiCarlo & Cox, 2007; Ito, Tamura, Fujita, & Tanaka, 1995; Lueschow, Miller, & Desimone, 1994; Marr & Nishihara, 1978; Tanaka, 1993). Adding an additional challenge, the features that define a target object may depend on learned associations, and can vary with task demands (e.g., when selecting a mango at the supermarket, the ideal appearance depends on your knowledge about ripe mangoes, as well as how quickly you plan to eat it). Thus, to accomplish target identification in real-world scenarios, representations of object target status must be invariant to low-level visual features, and able to integrate both visual and cognitive information. It is not yet known how these signals are generated in cortex, and how they give rise to behavioral outcomes.

Signals related to an object’s status as a target, or “match”, have been repeatedly reported in single units in feature and object selective visual areas. For instance, firing rate modulations in the middle temporal area (MT) signal when two directions of motion match (Lui & Pasternak, 2011), and changes in the gain and in the structure of V4 receptive fields selectively encode matches in shape and color (Bichot & Schall, 1999; Hayden & Gallant, 2013; Kosai, El-Shamayleh, Fyall, & Pasupathy, 2014). Many neurons in subregions of inferotemporal (IT) and entorhinal cortex also signal matches in the identity of complex objects (Miller & Desimone, 1994; Pagan, Urban, Wohl, & Rust, 2013; Woloszyn & Sheinberg, 2009). However, these match signals may not be invariant to low-level changes in object appearance, as most of these studies used targets that could be identified based on an exact match of retinal input patterns (Miller & Desimone, 1994; Pagan et al., 2013; Woloszyn & Sheinberg, 2009). Though basic invariance to many of these identity-preserving transformations has been demonstrated in subregions of the ventral visual stream (Anzellotti, Fairhall, & Caramazza, 2014; J. Erez, Cusack, Kendall, & Barense, 2016; Freiwald & Tsao, 2010; Tanaka, 1996), these past studies have all focused on representations of object identity itself, rather than investigating how target status is computed across changes in viewpoint. A few studies have addressed this gap by investigating the identification of relevant targets across changes in attributes like size and position (Lueschow et al., 1994; Roth & Rust, 2018). However, these studies focused exclusively on neural responses in sub-regions of the ventral visual stream and, importantly, did not examine target identification across changes in viewpoint. Because changes to an object’s viewpoint result in drastic differences in the 2D shape cast onto the retina, achieving target recognition across changes in viewpoint is likely to require more high-level, abstract search templates compared to when view invariance is not required (Biederman, 2001; Freiwald & Tsao, 2010; Riesenhuber & Poggio, 2000; Tarr, Williams, Hayward, & Gauthier, 1998).

While often considered a separate literature, studies focusing on the categorization of arbitrarily learned stimulus dimensions provide some indication that areas outside of ventral visual cortex may play a key role in generating viewpoint invariant target signals. For example, neurons in parietal and frontal cortex represent the membership of stimuli to learned categories, even when category membership is not predicted from visual similarity (Fitzgerald, Swaminathan, & Freedman, 2012; Freedman & Assad, 2016). Neurons in these regions may be selective to different directions of motion that belong to the same target category (Freedman & Assad, 2006), or even to different complex objects that belong to the same conceptual category (Freedman, Riesenhuber, Poggio, & Miller, 2001, 2003; Roy, Riesenhuber, Poggio, & Miller, 2010). Furthermore, in tasks that require identifying targets based on their membership to abstract categories, neurons in both prefrontal cortex (PFC) (Cromer, Roy, & Miller, 2010; Freedman et al., 2001, 2003; Roy et al., 2010) and premotor cortex (Cromer, Roy, Buschman, & Miller, 2011) encode objects’ target status. These findings suggest that when an object must be recognized based on abstract learned dimensions, target recognition may rely on computations that take place in areas outside of the ventral visual stream.

In addition to their ability to represent abstract visual information, regions of frontal and parietal cortex are also known to exhibit flexible coding, meaning their response properties are dependent on task context (Duncan, 2001; Miller, 2000; Miller & Cohen, 2001; Raposo, Kaufman, & Churchland, 2014). This makes them well suited for accomplishing target recognition under changing task demands. In particular, adaptive coding properties have been demonstrated in a set of parietal and frontal regions that are collectively referred to as the Multiple-Demand (MD) network (Duncan, 2010; Fedorenko, Duncan, & Kanwisher, 2013; Mitchell et al., 2016). Some sub-regions of the MD network have been previously shown to encode information about abstract task rules and response mappings (Harel, Kravitz, & Baker, 2014; Waskom, Kumaran, Gordon, Rissman, & Wagner, 2014; Woolgar, Thompson, Bor, & Duncan, 2011), as well as representations of relevant visual object properties (Y. Erez & Duncan, 2015; Jackson, Rich, Williams, & Woolgar, 2017; Vaziri-Pashkam & Xu, 2017). Though its role in invariant target recognition has not yet been tested, based on these coding properties, the MD network may play an important role in recognizing relevant objects during search, especially when successful search relies on flexibly updating the set of features that define the current target.

Given these gaps in the current knowledge about how invariant and flexible target representations are encoded to guide search behavior, we tested the hypothesis that regions of the MD network support performance during a target search task. To create a task scenario that required both invariance and flexibility, we generated a novel object stimulus set (Figure 1), in which 3D objects of multiple categories were rendered at multiple viewpoints. Subjects viewed sequentially presented objects while performing a one-back matching task in which they reported the status of each object as a match to the previous object in either identity (Identity Task) or viewpoint (Viewpoint Task), while ignoring similarity in the other dimension. Critically, this paradigm required subjects to form representations of abstract object dimensions that were invariant to the object’s retinal projection, and to flexibly attend to one object dimension depending on task instructions. We used MVPA on single-trial voxel activation patterns to decode the status of each image as a match in each dimension and compared decoding performance for matches in the task-relevant dimension versus the task-irrelevant dimension. Moreover, we compared the strength of decoding on correct and incorrect trials to assess the relative association of modulations in different cortical regions with behavior. Our findings suggest that, while ventral visual cortex exhibits some sensitivity to an object’s status as a target, regions of the MD network encode robust, invariant target representations that are sensitive to changes in task demands and that are selectively linked with behavioral performance.

**Figure 1.**
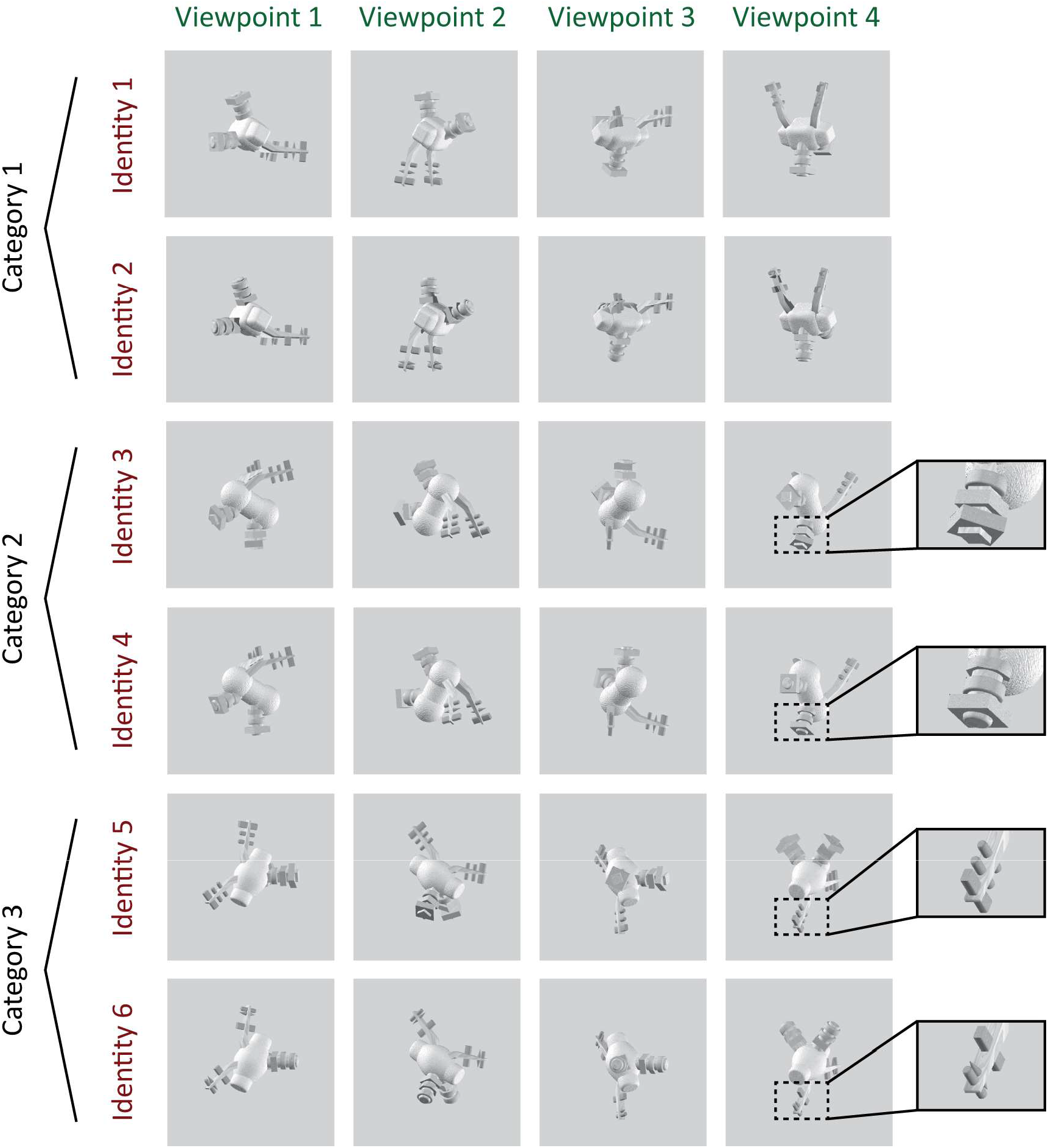
Example set of images shown to a subject during scanning, consisting of six unique object identities, each rendered at four viewpoints. Subjects were instructed either to match the exact identity of the object irrespective of viewpoint (shown in rows of the matrix) or to match the viewpoint of the object irrespective of identity (columns of the matrix). The six identities comprised two exemplars in each of three categories, with categories defined by overall body shape, and exemplars defined by details of the peripheral features (see inset panels for examples of differentiating features). Object viewpoint was generated in an arbitrarily-defined coordinate system so that low-level visual features had a minimal contribution to the viewpoint matching task (see *Methods* for details). Two complete sets of novel objects (Set A and Set B) were generated, with half the subjects (5/10) viewing set A, and half viewing Set B. These images are from Object Set B, see Supplementary Figure 1 for examples of Object Set A.

## Results

### Behavioral Performance

Subjects (n=10) performed alternating runs of the Identity Task and the Viewpoint Task while undergoing fMRI. Runs were always presented in matched pairs, so that the object sequence and visual stimulation was identical between runs of the Identity Task and Viewpoint Task. We found no significant difference in performance (Figure 2A; d’ for Identity: 1.47 ± 0.24, d’ for Viewpoint: 1.81 ± 0.34; paired two-tailed t-test, p=0.2054), and no significance difference in response times (Figure 2B, RT for Identity: 1.46 ± 0.05 s, RT for Viewpoint: 1.35 ± 0.03 s; paired two-tailed t-test, p=0.1313) on the two tasks across subjects. Performance and response time for each task also did not differ as a function of the object set subjects had been assigned to (d’ for Identity, Set A: 1.50 ± 0.22, d’ for Identity, Set B: 1.43 ± 0.27, p=0.8930; d’ for Viewpoint, Set A: 1.76 ± 0.30, d’ for Viewpoint, Set B: 1.86 ± 0.42, p=0.9024; RT for Identity, Set A: 1.38 ± 0.03 s, RT for Identity, Set B: 1.54 ± 0.06 s, p=0.1486; RT for Viewpoint, Set A: 1.36 ± 0.03, RT for Viewpoint, Set B: 1.35 ± 0.04, p=0.8758; all are two-tailed t-tests).

**Figure 2.**
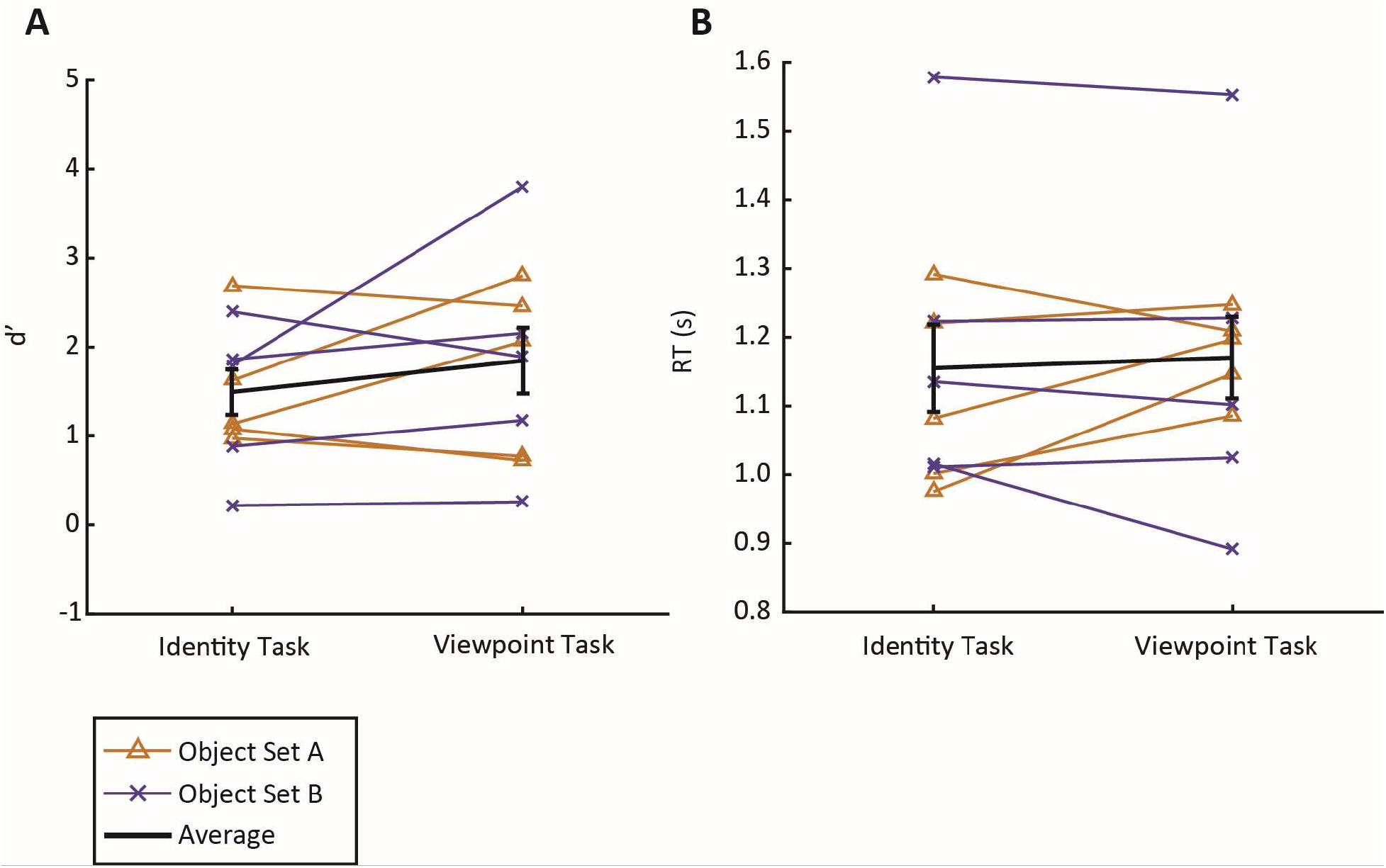
Behavioral performance (d’) and response time (RT) were similar across tasks and stimulus sets. Each line represents performance of a single subject, averaged over runs of each task. Error bars indicate ± 1 SEM.

### Univariate BOLD Signal does Not Show Match Suppression

First, we examined whether the status of each image as a match in the task-relevant dimension (e.g., identity match status during the Identity Task; viewpoint match status during the Viewpoint Task) was reflected in a change in the mean amplitude of BOLD response in any visual area. Based on previous fMRI studies (Grill-Spector et al., 1999; Henson, 2003) and electrophysiology studies (Meyer & Rust, 2018; Miller, Li, & Desimone, 1991), we predicted that the repetition of object identity or viewpoint might result in response suppression (often referred to as repetition suppression). In all but one of the 14 ROIs we examined (Figure 3), we found no significant difference in the mean signal amplitude between matches and non-matches. This indicates a lack of evidence for repetition suppression based on match status at the level of the univariate BOLD signal. In the one ROI showing a significant effect (AI-FO), mean signal was actually higher on match trials than non-match trials. We suspect that the absence of repetition suppression is due to differences in task demands between our experiment and previous work, particularly the fact that identity and viewpoint matches were task-relevant in our paradigm (see *Discussion* for details). Note that this analysis was performed before the data were de-meaned; in all subsequent multivariate analyses, we de-meaned the data so that each single trial voxel activation pattern was centered at zero and any small univariate effects could not contribute to decoding performance (see *Methods*).

**Figure 3.**
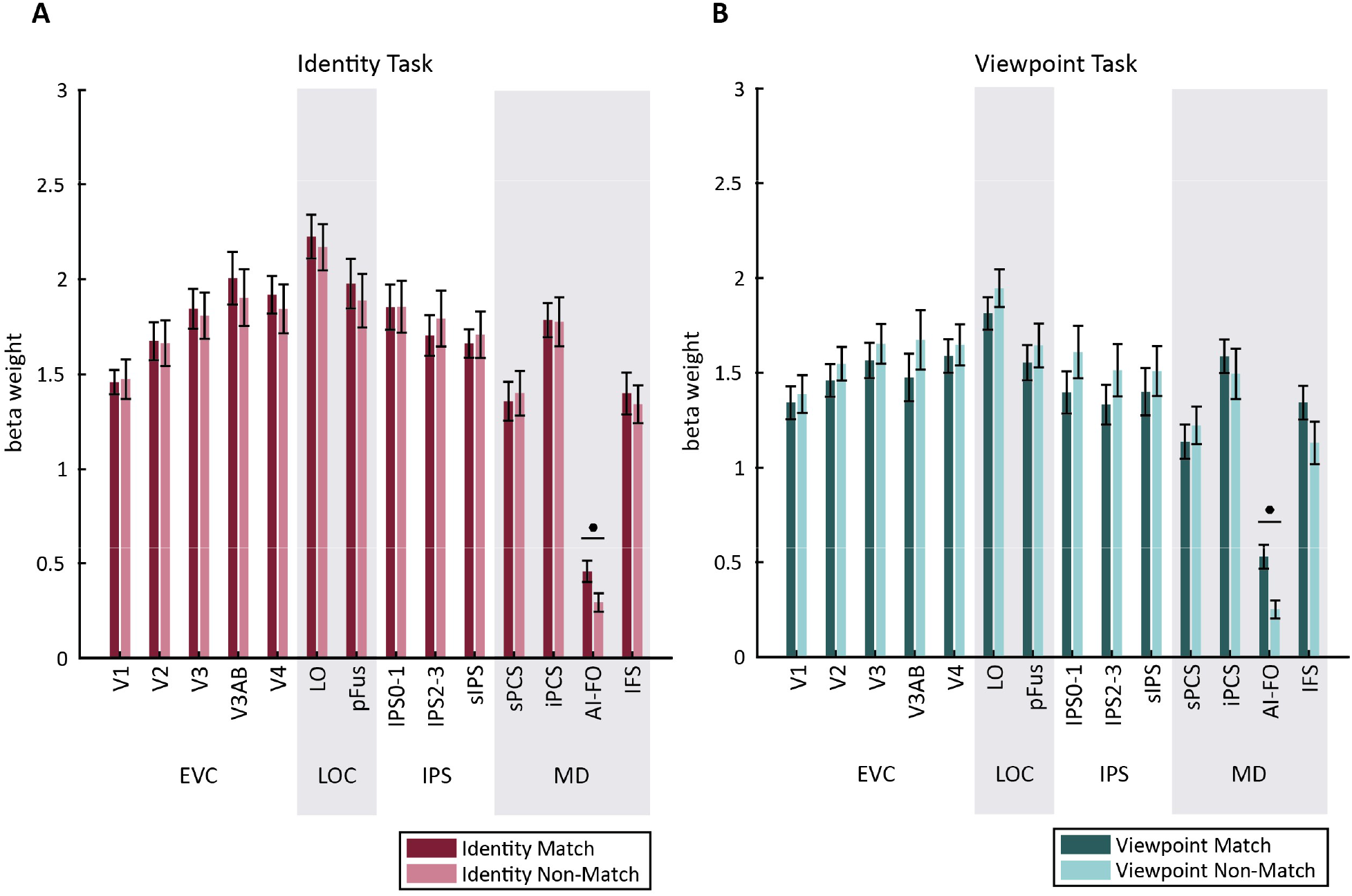
Match status in the current task cannot be determined solely from mean signal change. Mean beta weights are plotted on the Y axis, and individual ROIs are plotted on the X axis. In all fMRI analyses, we used 14 ROIs: early visual cortex (V1, V2, V3, V3AB, and V4), the lateral occipital complex (LO, pFus), the intraparietal sulcus (IPS0-1, IPS2-3, superior IPS/sIPS), and the multiple-demand network (superior precentral sulcus/sPCS, inferior precentral sulcus/iPCS, anterior insula-frontal operculum/AI-FO, and inferior frontal sulcus/IFS), all of which were defined using functional localizers (see *Methods*). ROIs are organized into four groups for convenience. Note that though we have visually separated them in all figures, we consider IPS to be a part of the MD network. **(a)** In the Identity Task, in all but one ROI, univariate activation (mean beta weight) did not differ between trials according to their status as an identity match. **(b)** Similarly, in the Viewpoint Task, univariate activation in most ROIs did not reflect viewpoint match status. Match and non-match trials were compared using paired t-tests, p-values were FDR corrected over all conditions; solid circles indicate significance at q=0.01. Error bars indicate ± 1 SEM.

### Multivariate Activation Patterns Reflect Task-Relevant Match Status

Next, we examined how voxel activation patterns in each ROI reflected the status of a stimulus as a match in viewpoint and identity, and how representations of viewpoint and identity match status were influenced by the task-relevance of each dimension. During each task, a correct behavioral response depended on the object being a match to the previous object in the relevant dimension (identity or viewpoint), while match status in the other dimension was irrelevant. Therefore, we expected that information about the match status of an object in each dimension would be more strongly represented when that dimension was task-relevant.

Indeed, status as a match along the task-relevant dimension, measured by classifier performance (d’), was represented widely within the ROIs we examined, while the irrelevant match was not represented at an above-chance level in any ROI (Figure 4). Information about the task-relevant match increased along a posterior-to-anterior axis, such that match status was represented most strongly in MD and IPS ROIs, but was comparatively weaker in early visual cortex and the LOC. Relevant match decoding performance was above chance for all MD ROIs for both the identity and viewpoint task, and was also above chance for LO, V3AB, and V2 for both tasks. Decoding performance was above chance in V1, V4, and pFus for the identity task only.

**Figure 4.**
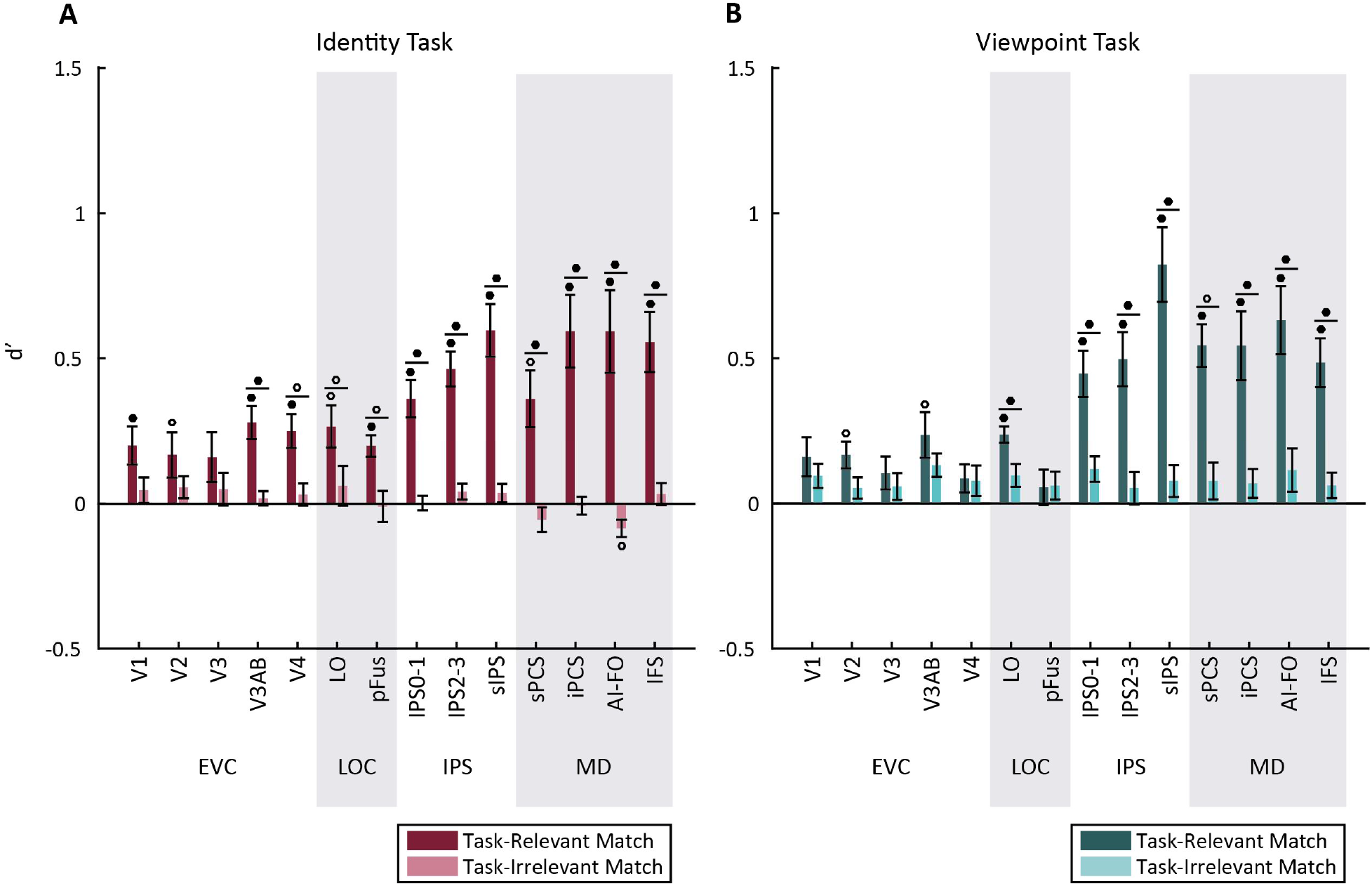
Task-relevant matches are represented more strongly than task-irrelevant matches. A linear classifier was trained to discriminate between voxel activation patterns measured during each task, according to whether the viewed image was a match in the task-relevant and the task-irrelevant dimension. Classifier performance (d’) is plotted on the Y axis. Circles above individual bars indicate above-chance classification performance (test against zero); circles above pairs of bars indicate significant differences between bars (paired t-test). P-values were FDR corrected over all conditions; open circles indicate significance at q=0.05, solid circles indicate significance at q=0.01. Error bars indicate ± 1 SEM.

The general pattern of decoding performance was similar across tasks, though there was a trend toward higher relevant match decoding performance in IPS for the viewpoint task than the identity task. There was also an opposite trend in V4 and pFus for higher relevant match decoding during the identity task than the viewpoint task. A three-way repeated measures ANOVA with factors of Task, ROI and Relevance revealed a main effect of Relevance (F(1,9) = 46.348, p=10^−4^), a main effect of ROI (F(13,117) = 9.918, p<10^−4^), and a Relevance x ROI interaction (F(13,117) = 13.976, p<10^−4^), but no main effect of Task or Task-related interactions were observed. We further investigated the ROI x Relevance interaction using paired t-tests to compare decoding of the relevant and irrelevant dimensions, and found that the effect of relevance was significant for all MD regions in both tasks, LO in both tasks, as well as V3AB, V4, and pFus for the identity task only (Figure 4).

### Control Analyses for Visually-Driven Match Representations

To perform the identity and the viewpoint tasks, subjects were required to use representations of object identity and viewpoint that were largely invariant to shape. However, given a limited number of trials, each task could, in principle, be solved based only on shape similarity between the previous and current objects. In the viewpoint task, since subtle differences between exemplars were not task-relevant, the group of trials that could be solved based only on shape similarity included all trials that were matches in both category and viewpoint. In the identity task, the group of trials that could be solved based on shape similarity included all trials that were an exact match to the previous image in category, exemplar, and viewpoint. Therefore, it is possible that some of the match status classification we observed (Figure 4) was driven by the detection of shape similarity. To test for this possibility, we removed all trials that were a match in both category and viewpoint, leaving a set of trials in which the shape similarity was entirely uninformative about match status. We then performed classification on this reduced dataset as before.

For the viewpoint task, we now found an important difference between visual and MD regions: While decoding of viewpoint matches remained above chance in all MD and IPS regions, it dropped to chance in LOC and early visual cortex (Figure 5B). Thus, while early visual and LOC representations of viewpoint match status appeared to rely on low-level shape similarity, MD regions encoded viewpoint match status even when shape similarity could not be used to define a match.

**Figure 5.**
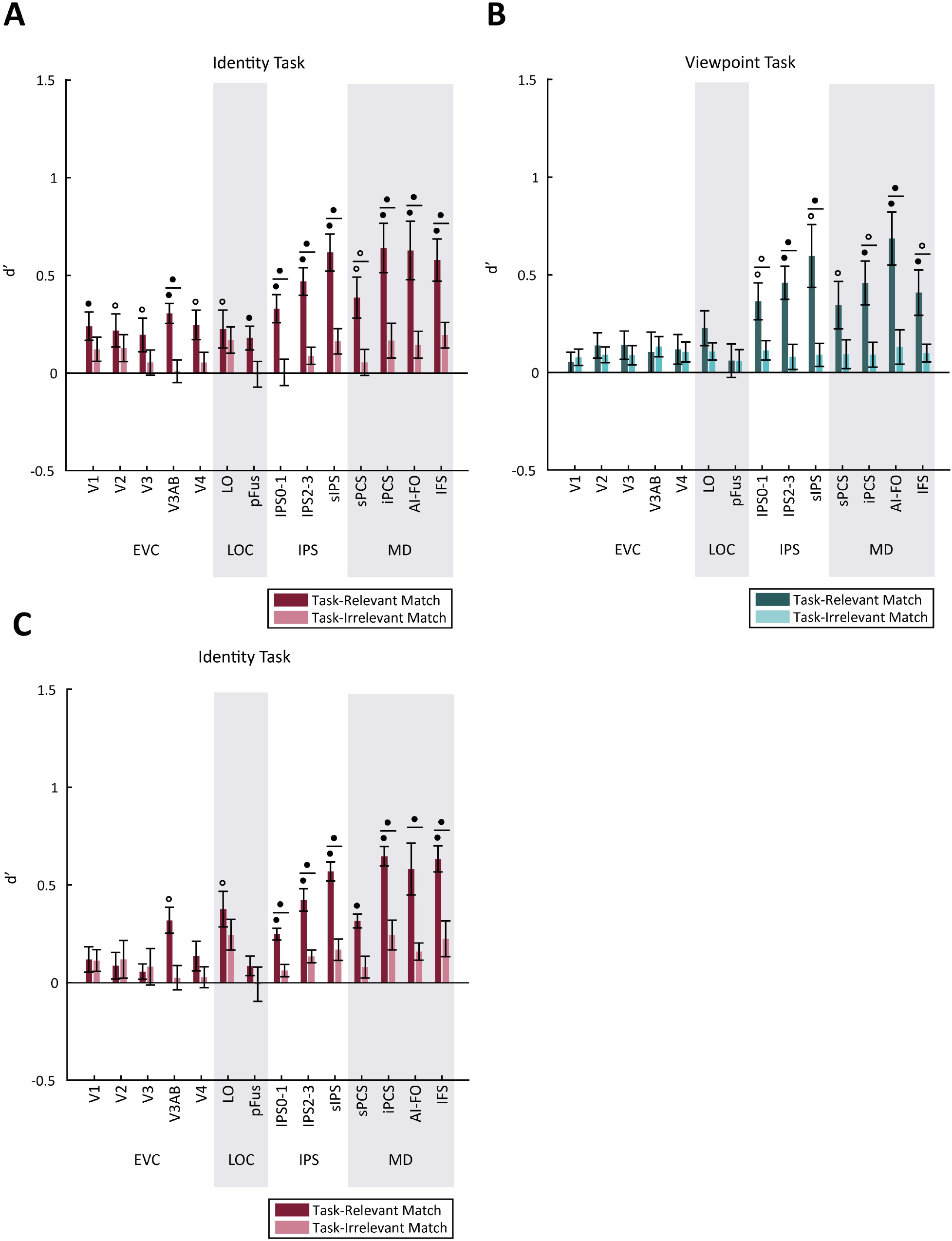
Control analyses related to Figure 4: viewpoint and identity match information in MD regions is not driven by low-level image statistics. **(a and b)** To address the possibility that match status could have been inferred from low-level visual properties, we removed all trials in which an object had a high degree of shape similarity to the previous object, and repeated the analyses of Figure 4. **(a)** Identity match information remained above chance in all regions. **(b)** Viewpoint match information dropped to chance in the early visual cortex and LOC, but remained above chance in MD regions. **(c)** After identifying that the pairwise similarity between images in Object Set A was informative about identity match status, we re-analyzed the identity task data using only the subjects shown Object Set B. Identity match classification in early visual and ventral visual cortex drops below significance when subjects in Object Set A are removed, but remains above chance in MD regions. P-values were computed at the subject level over these 5 subjects and FDR corrected across ROIs; open circles indicate significance at q=0.05, solid circles indicate significance at q=0.01. Error bars indicate ± 1 SEM.

During the identity task, however, even when shape similarity could not be used to define match status, decoding of identity match status remained above-chance in all ROIs examined (Figure 5A). The observation of above-chance identity match decoding in early visual areas was surprising given that these areas are not expected to encode abstract, viewpoint independent representations of object identity. Therefore, we wondered whether either of our stimulus sets made it possible for identity match status to be inferred based on shape similarity. Though we had already removed all trials in which the previous and current images were highly similar in shape, the remaining set of images may have had some shared features that supported match classification. Indeed, when we assessed the pixel-wise similarity between images (see *Methods*) belonging to each stimulus set, we found that in 4 of the subjects assigned to Object Set A, pixel-wise similarity between images was significantly predictive of identity match status (Supplementary Figure 2). We thus hypothesized that the above-chance decoding of identity match status observed in early visual areas might be driven by this group of subjects.

In line with this prediction, when we re-analyzed decoding performance using only subjects from Set B, identity match decoding was no longer significant in V1, V2, V3, V4, and pFus, suggesting that the above-chance decoding accuracy for identity match status in these ROIs had been driven at least in part by low-level image features (Figure 5C). Despite this drop, identity match decoding performance remained above chance in all MD and IPS ROIs. Therefore, when low-level image features were uninformative about identity match status (Object Set B), MD and IPS regions still encoded robust representations of identity match status, while early visual cortex did not.

### Behavioral Performance is Closely Linked to Activation Patterns in the MD Network

Having established that several ROIs, including all MD network ROIs, represent the status of an object as a match in the task-relevant dimension, we next sought to determine whether these representations were related to task performance. To answer this question, we used the normalized Euclidean distance to calculate a continuous measure of the classifier’s evidence at predicting the correct label for each trial in the test set (see *Methods*). We then calculated the mean classifier evidence for all correct and incorrect trials. Because the previous analysis (Figure 4) indicated no significant effect of task on relevant match decoding, we combined all trials across the identity and viewpoint tasks for this analysis (Figure 6).

**Figure 6.**
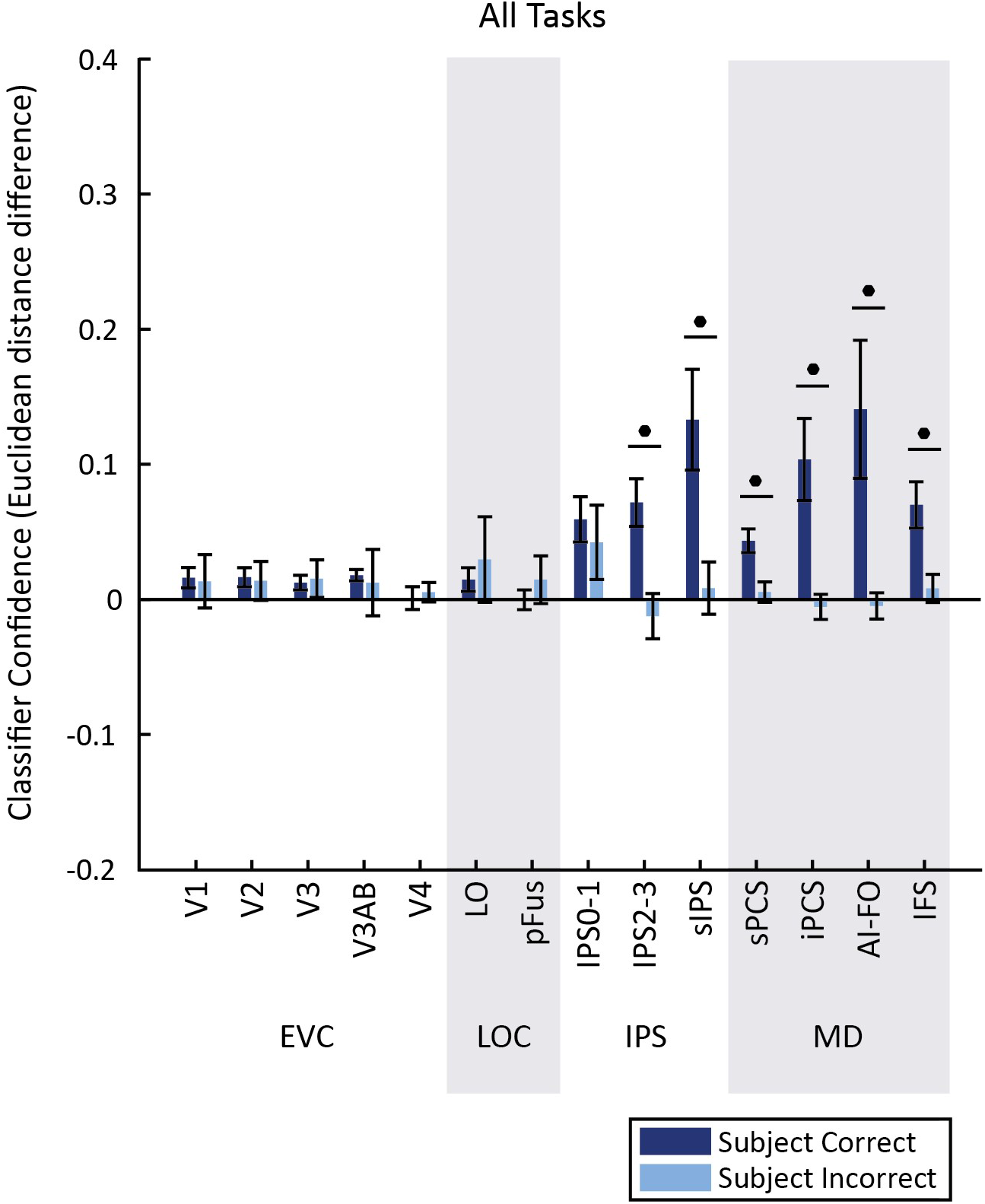
Classifier evidence is associated with task performance. Trials were classified according to their status as a match in the task-relevant feature, and the normalized Euclidean distance was used as a metric of evidence in favor of the correct classification (distance to incorrect label – distance to correct label, plotted on the Y axis). Circles above pairs of bars indicate a significant difference between correct and incorrect trials (paired t-test). Solid circles indicate significance at q=0.01 (FDR corrected). Error bars indicate ± 1 SEM.

In all regions of the MD network and IPS, we found that classifier evidence was significantly higher on correct than incorrect trials. In contrast, we found no significant effect of behavior on classifier evidence in early visual cortex or the LOC. Thus, representations in IPS and MD ROIs, but not any of the other regions that we evaluated, were significantly associated with task performance.

## Discussion

Here we used fMRI and pattern classification methods to investigate the role of different brain areas in signaling task-relevant matches across identity preserving transformations. Specifically, subjects performed a task that required them to identify matches in either the identity or the viewpoint of novel objects. Consistent with previous work, we found that areas of ventral visual cortex represent information about the status of an object as a relevant match (Figure 4; Lueschow et al., 1994; Miller & Desimone, 1994; Pagan et al., 2013; Woloszyn & Sheinberg, 2009). However, our results also suggest a key role for the MD network in match identification, with MD match representations showing specificity to task-relevant dimensions (Figure 4), as well as invariance to low-level visual features (Figure 5). Additionally, there was a significant link between task performance and the strength of MD match representations (Figure 6). In contrast, in early visual and ventral object-selective cortex, we found comparatively weaker evidence for representation of match status, and no association with task performance. These results suggest that MD regions play a key role in computing flexible and abstract target representations and have a gating influence on task performance.

In contrast to our MVPA results, univariate signal amplitude in almost all ROIs was not significantly modulated by match status. This finding differs from many past fMRI studies (Grill-Spector et al., 1999; Henson, 2003; Turk-Browne, Yi, Leber, & Chun, 2007) which have observed response suppression as a result of object repetition (or “repetition suppression”). One explanation for this divergence in findings is that in our task, the repetition of identity and viewpoint is task-relevant, meaning that repetition related signals are mixed with signals related to task performance. Many electrophysiology studies have found that when an object’s match status is task-relevant, neural response modulations are heterogeneous, including both enhancement and suppression (Engel & Wang, 2011; Lui & Pasternak, 2011; Miller & Desimone, 1994; Pagan et al., 2013; Roth & Rust, 2018). This type of signal would be detectable using multivariate decoding methods, but in univariate analyses may be obscured by averaging across all voxels in an ROI (Serences & Saproo, 2012). Therefore, our data are consistent with the interpretation that match representations are not an automatic by-product of stimulus repetition, but are linked to the task-relevance of each stimulus.

Representations of identity and viewpoint match status were weaker in early visual and ventral ROIs compared to ROIs in the MD network. Moreover, several control analyses suggest that the representations in some of these occipital and ventral regions were driven primarily by low-level image statistics, as opposed to an object’s status as a match in the task-relevant dimension. First, after removing all trials in which the current and previous objects were matches in both category and viewpoint, we found that viewpoint match decoding during the viewpoint task dropped to chance in all ROIs except for those in the MD network. Additionally, a post-hoc image-based analysis revealed that one of our stimulus sets had a systematic bias in pixel-similarity, such that the degree of image similarity between each pair of objects was informative about the status of the pair as an identity match. When we repeated our decoding analysis using only subjects who had seen the non-biased set, we found that identity match decoding dropped to chance in V1, V2, V3, V4, and pFus, though it remained above chance in V3AB and LO. Together, these findings suggest that low-level visual features were at least partially responsible for the viewpoint and identity match representations we initially observed in early visual and ventral visual areas.

While match representations in early visual cortex and the LOC were weaker than those measured in MD regions, we did observe a consistent modulation of these representations by task, such that the relevant match was generally represented more strongly than the irrelevant match. This modulation was significant in LO during both tasks, and in V3, V3AB, and pFus during the identity task (Figure 4). One interpretation for the enhanced representation of relevant match status is the presence of non-specific reverberating activity related to relevant match representations computed at higher levels of processing. This is supported by the observation that the match representations in early and ventral ROIs were not, in general, significantly associated with behavior. Alternatively, the observation of task-related modulations suggests a role for top-down feedback in determining the content of visual representations in early visual and ventral object selective cortex. This feedback could act to enhance representations of visual properties that are informative for computing match status, which would result in an enhancement of signals related to repetition of these properties

Overall, our results suggest that relevant match representations in MD regions were significantly associated with behavior, more so than in regions of occipital or ventral visual cortex. Moreover, representations of match status in occipital and ventral ROIs may have been driven primarily by low-level shape information, while representations in the MD network were largely unaffected by the removal of such low-level information. Together, these findings suggest that frontoparietal MD regions tend to represent objects in a more abstract, task-dependent way, while ventral regions tend to be more visually driven and shape-dependent.

This view is supported by several previous studies. A recent fMRI study found that abstract face-identity information is represented more strongly in IPS than in LO and the FFA (Jeong & Xu, 2016). Another study found that during the delay period of a working memory task, object information could be decoded from PFC only when the task was nonvisual, and from the posterior fusiform area only when the task was visual, supporting a dissociation between these regions in representing abstract and visual object information respectively (Lee, Kravitz, & Baker, 2013). In another study, primates were trained to perform categorization on a set of morphed animal stimuli designed so that category membership was not predicted by visual similarity of individual images (Freedman et al., 2003). They found that the visual responses of IT cortex neurons could be explained by selectivity for individual images, while responses of PFC neurons showed category selectivity above that expected from a purely stimulus-selective population. Furthermore, when monkeys performed a category matching task, the amount of match selectivity observed in ITC was lower than a previous study which had used an identity matching paradigm, suggesting that ITC might be more involved in comparison of identities than categories (Miller & Desimone, 1994; Miller et al., 1991). Overall, our findings provide additional support for the conclusion that frontoparietal cortex represents abstract object dimensions and task-relevant information, while ventral regions are more involved in representing the details of currently-viewed images.

That said, our results do not rule out the possibility of abstract object information in any portion of the ventral stream. Many past studies have demonstrated viewpoint-invariant object information in human and primate ventral visual cortex (Anzellotti et al., 2014; J. Erez et al., 2016; Freiwald & Tsao, 2010; Tanaka, 1996). In agreement with this role in object processing, our univariate analysis (Figure 3) showed the highest mean signal amplitude in area LO during both the identity and viewpoint tasks. Despite this, our multivariate decoding analysis failed to detect an association with behavior in any ventral ROI. One explanation for this finding is that target information was present at a level of granularity below what is detectable with our methods, or that the signal to noise regime in ventral object selective cortex prevented us from detecting abstract object representations (Dubois, de Berker, & Tsao, 2015). Furthermore, it is possible that linearly-decodable information about abstract object properties may be more readily detectable at later stages of the ventral visual stream, such as perirhinal cortex, as compared to LO and pFus (J. Erez et al., 2016; Pagan et al., 2013). Future studies will be needed to determine the extent to which abstract matches in identity and viewpoint may be represented within ventral visual cortex.

Finally, a novel finding of this study is that MD regions contain representations of matches in viewpoint across objects, in addition to matches in identity. This suggests that beyond just representing abstract object identity, MD network regions may be capable of representing object viewpoint in a format that is invariant to the identity and shape of objects. A recent modeling study found that at ascending positions in the ventral visual stream, information about “category-orthogonal” object properties such as viewpoint, position, and size increases in parallel to information about object category and identity, and is extracted by the same computational mechanisms that compute invariant object identity representations (Hong, Yamins, Majaj, & Dicarlo, 2016). Our finding of both viewpoint-invariant identity information and identity-invariant viewpoint information in MD regions supports the idea that these two orthogonal types of information may be computed by related mechanisms.

In conclusion, our findings support a role of MD network regions in the comparison of high-level object dimensions, such as viewpoint and identity, between remembered and viewed visual objects. The match representations in these regions are modulated by task-relevance, and are associated with subject behavior on a trial-by-trial basis. In contrast, early visual and ventral object selective cortex contained weaker evidence for match representations, with some modulation by task, consistent with a role for top-down feedback in enhancing representations of relevant information at the earliest levels of processing.

## Methods

### Participants

10 subjects (3 male) between the ages of 20 and 34 were recruited from the UCSD community (mean age 24.7 ± 4.7 years), having normal or corrected-to-normal vision. This sample size was determined before starting data collection, based on sample sizes used by past experiments with similar methodology. All participants provided written informed consent in accordance with the local Institutional Review Board at UCSD. Each subject performed a behavioral training session lasting approximately 1 hour, followed by 1 or 2 scan sessions, each lasting approximately 2 hours. Participants were compensated at a rate of $10/hour for behavioral training and $20/hour for the scanning sessions.

### Novel Object Sets and One-Back Tasks

We generated two novel object stimulus sets for this study (see Figure 1 and Supplementary Figure 1), which had several advantages. First, designing the objects in-house allowed us to select body configurations that would have drastically different shapes when viewed from different viewpoints, making it challenging to use the 2D shape of an object to determine its identity. Second, we were able to define object viewpoint in an arbitrarily defined coordinate system, so that it was impossible for subjects to determine object viewpoint based on 2D object shape. This coordinate system was defined as follows. For each object category, we defined a single viewpoint of the object as its “frontal” viewpoint, or a rotation of [0°,0°] about the Y and Z axes. We assigned an arbitrary viewpoint as “frontal” so that this canonical viewpoint was not systematically predictable from the overall axis of elongation of the main body. As a result, the frontal viewpoints of the objects in different categories differed dramatically in the shape they produced when projected onto a 2D plane. This ensured that subjects formed a representation of viewpoint as an [X°, Y°] pair that was largely invariant to 2D shape.

Prior to scanning, subjects participated in a self-guided exercise to familiarize them with the novel object viewpoints. During the first half of this training, subjects viewed a 3D model of each of the three novel object categories, and were able to rotate the model around two axes using the arrow keys on a keyboard (each key press gave a rotation of 30 degrees about either the Y or Z axis). During this entire exercise, the angular position of the object, expressed using the format “Y rotation = m degrees, X rotation = n degrees”, was displayed at the top of the screen. Subjects were encouraged to use the angular coordinates to learn how each view of the object was defined relative to the frontal ([0°,0°]) position. During the second half of this training, subjects were presented with images of all three categories simultaneously, at matching viewpoints, and encouraged to study how the same viewpoint was defined across categories. Subjects performed both parts of this training at least once, and were allowed to return to it as many times as they wished. On average subjects spent approximately 20 minutes on the self-guided training.

All objects were generated and rendered using Strata 3D CX software (version 7.6; Santa Clara, UT). To ensure that our results were not an artifact of any idiosyncratic aspect of one stimulus set, we generated two unique sets of objects, and assigned half of our subjects (five out of 10) to Object Set A, and half to Object Set B (since we did not observe a difference in performance between the two stimulus sets for either task, we combined our analyses across all subjects). Each stimulus set comprised 3 categories of objects, each with 36 total exemplars. Objects within a category shared a common body plan, including the shape of the main body and the configuration of peripheral features around the body. All objects in a given set had the same peripheral features, which differed across categories only in how they were attached to the body. There were two peripheral feature types, each of which appeared twice on each object. Pairs of matching features were attached symmetrically to the body, making the overall objects bilaterally symmetric. Exemplars in each category were differentiated by small variations in the details of peripheral features, such as the size or shape of a spike. There were 6 variants for each feature pair, giving 36 total exemplars of each category. Feature details were always matched within each peripheral feature pair, assuring that even when one feature in a pair was occluded at a particular viewpoint, the details could always be discerned from the other feature in the pair.

Though there were 36 exemplars per category in each full object set, each subject only viewed two exemplars of each category during scanning. The selection of these two exemplars allowed us to manipulate the difficulty of the task for each subject, by selecting exemplar pairs that were ~70% confusable. To select these exemplars, each subject participated in a behavioral experiment before the scan. This task was identical to the one-back Identity task used during scanning, but it employed a limited stimulus set in which objects were drawn from a single category and only differed in one of the two peripheral feature pairs. Subjects were told at the start of this task which of the feature pairs was relevant for task performance, and that changes in viewpoint were task-irrelevant. Each subject completed six runs of this training task, with each object category and each feature type serving as the main focus for each run. In analyzing the confusability of variants of the two feature types, we collapsed across object category. We then selected, for each feature type, three pairs of variants that were confused approximately 70% of the time. These three pairs were then randomly assigned to the three object categories, giving two total exemplars per object category. Exemplars in the same category always differed in both feature types, but exemplars in different categories could have the same variant of one or both feature types.

While in the scanner, each subject performed two different one-back tasks (Identity Task and Viewpoint Task). Task runs always occurred in pairs with the identity task followed by the viewpoint task, and each subject performed between 8 and 11 pairs of runs in total. An identical sequence of visual stimuli was presented on both runs in each pair, so that visual stimulation was perfectly matched between conditions.

Each run consisted of 48 trials, and each trial consisted of a single image presentation. During each run, six different objects were shown (two exemplars in each of three categories), from four different viewpoints; each image was shown twice. The sequence of image presentations was generated using a pseudo-randomization procedure, with the constraint that the probability of the current stimulus being from the same category as the previous stimulus (within-category trials) was approximately 0.50. This constraint was used to more closely equate the difficulty of the two tasks, as the viewpoint task was more difficult to solve on across-category trials, and the identity task was more difficult to solve on within-category trials. This resulted in a probability of 0.23 of any trial being a match in either viewpoint or identity, and a probability of 0.04 of a match in both dimensions.

In the Identity Task, subjects responded to each image based on whether it matched the identity of the immediately preceding image. Identity matches had to be both from the same category and the same exemplar, but did not have to match in viewpoint. In the Viewpoint Task, subjects responded based on whether the current image matched the viewpoint of the immediately preceding image, while both category and exemplar status was irrelevant. For instance, an object from category A presented at a viewpoint of [30°, 150°] would be a viewpoint match to an object from category B presented at [30°, 150°].

Before scanning, subjects performed several practice runs of both the Identity Task and the Viewpoint Task. During these runs, the four object viewpoints that were used during scanning were never used. After each Viewpoint Task practice run, subjects could return to the self-guided viewpoint training, and repeated as many iterations of self-guided training and practice runs as were necessary to reach 70% performance.

Immediately prior to each scanning session, subjects performed another short self-guided training exercise (~5 minutes), in which they were shown examples of the exact images they would see during scanning. First, they were presented with the two exemplars in each object category, side-by-side, and allowed to freely rotate the models using the arrow keys and compare the appearance of the two exemplars from many viewpoints. Next, they were presented with images from each category, side-by-side, at each of the four viewpoints they would see during the task, and encouraged to use this information to prepare for the viewpoint task.

In both tasks, subjects responded to every image by pressing a button using either their index finger (“1”) or their middle finger (“2”), depending on the current response mapping rule. Response mapping rules were counterbalanced within each subject, so that on half the runs the subject responded with 1 for “match” and 2 for “non-match”, and on the other half of runs they responded with 1 for “non-match” and 2 for “match”. The purpose of these different response mapping rules was to ensure that match-related information was not confounded with motor response.

### Magnetic Resonance Imaging

All MRI scanning was performed on a General Electric (GE) Discovery MR750 3.0T research-dedicated scanner at the UC San Diego Keck Center for Functional Magnetic Resonance Imaging (CFMRI). Functional echo-planar imaging (EPI) data were acquired using a Nova Medical 32-channel head coil (NMSC075–32-3GE-MR750) and the Stanford Simultaneous Multi-Slice (SMS) EPI sequence (MUX EPI), with a multiband factor of 8 and 9 axial slices per band (total slices = 72; 2 mm^3^ isotropic; 0 mm gap; matrix = 104 × 104; FOV = 20.8 cm; TR/TE = 800/35 ms; flip angle = 52°; inplane acceleration = 1). Image reconstruction procedures and un-aliasing procedures were performed on local servers using reconstruction code from CNI (Center for Neural Imaging at Stanford). The initial 16 TRs collected at sequence onset served as reference images required for the transformation from k-space to the image space. Two short (17 s) “topup” datasets were collected during each session, using forward and reverse phase-encoding directions. These images were used to estimate susceptibility-induced off-resonance fields (Andersson, Skare, & Ashburner, 2003) and correct signal distortion in EPI sequences, using FSL topup functionality (Jenkinson, Beckmann, Behrens, Woolrich, & Smith, 2012).

During each functional session, we also acquired an accelerated anatomical scan using parallel imaging (GE ASSET on a FSPGR T1-weighted sequence; 1×1×1 mm^3^ voxel size; 8136 ms TR; 3172 ms TE; 8° flip angle; 172 slices; 1 mm slice gap; 256×192 cm matrix size) using the same 32-channel head coil. We also acquired an additional high-resolution anatomical scan (1×1×1 mm^3^ voxel size; 8136 ms TR; 3172 ms TE; 8° flip angle; 172 slices; 1 mm slice gap; 256×192 cm matrix size) during a separate retinotopic mapping session using an Invivo 8-channel head coil. This scan produced higher quality contrast between grey and white matter and was used for segmentation, flattening, and visualizing retinotopic mapping data.

In addition to the multiband scan protocol described above, several subjects participated in retinotopic mapping experiments using a different scan protocol, previously reported (Sprague & Serences, 2013). Where appropriate, the data used to generate retinotopic maps (see *Retinotopic Mapping*) was combined across these sessions.

### Pre-Processing

First, the structural scan from each session was processed in BrainVoyager 2.6.1 to align the anatomical and the functional datasets. Automatic algorithms were used to adjust the structural image intensity to correct for inhomogeneities, as well as remove the head and skull tissue. Structural scans were then aligned to the AC-PC plane using manual landmark identification. Finally, an automatic registration algorithm was used to align the structural scan to the high-definition structural scan collected during each subject’s retinotopic mapping session.

Next, each functional run was aligned to the same-session structural scan. We then performed slice-time correction, affine motion correction, and temporal high-pass filtering to remove slow signal drifts over the course of each session in BrainVoyager 2.6.1. These data were spatially transformed into Talairach space. Finally, the BOLD signal in each voxel was z-transformed within each run. Single-trial activation estimates (beta weights), which were used for MVPA analyses, were obtained by solving a general linear model (GLM) with a design matrix created by convolving trial events with a canonical two-gamma HRF (peak at 5 s, undershoot peak at 15 s, response undershoot ratio 6, response dispersion 1, undershoot dispersion 1). Throughout this paper, the same HRF parameters were used for all GLM analyses.

### Retinotopic Mapping

We followed previously published retinotopic mapping protocols to define the ROIs reported here (Jerde & Curtis, 2013; Swisher, Halko, Merabet, McMains, & Somers, 2007; Wandell, Dumoulin, & Brewer, 2007; Winawer & Witthoft, 2015). Subjects performed mapping runs in which they viewed either rotating wedge (10 cycles, 36 s/cycle), expanding ring (10 cycles, 36 s/cycle), or bowtie (8 cycles, 40 s/cycle) stimuli. To increase the quality of data from parietal regions, subjects performed a covert attention task on the rotating wedge stimulus, which required them to detect contrast dimming events in a row of the checkerboard (mean accuracy = 61.8 ± 13.9%). This stimulus was limited to 22° by 22° FOV.

### Multiple-Demand (MD) Localizer

To define ROIs in the MD network, we used a functional localizer to identify voxels whose BOLD response was significantly modulated by the load of a spatial working memory task (Duncan, 2010; Fedorenko et al., 2013). Subjects performed 1–2 runs of this task during each functional scanning session. During each trial of this task, subjects were first presented with an empty rectangular grid comprising either 8 or 16 squares. Half of the squares in the grid were then highlighted one at a time, and subjects were required to remember the locations of the highlighted squares. Subjects were then shown a probe grid and asked to report whether the highlighted squares matched the remembered locations. Runs were divided into blocks with either high or low load. Performance was significantly poorer on high load blocks (mean d’ for low load = 2.48 ± 0.28, mean d’ for high load = 1.04 ± 0.21, p < 0.001; paired two-tailed t-test).

We used the data from these runs to generate a statistical parametric map for each subject, which expressed the degree to which each voxel showed elevated BOLD signal for high versus low load working memory blocks. We defined a GLM with a regressor for each block type, and solved for the beta coefficients corresponding to each load condition. Coefficients were then entered into a one-tailed, repeated-measures t-test against a distribution with a mean of 0 (FDR corrected q=0.05). This resulted in a single mask of load-selective voxels for each subject.

To subdivide this mask into the MD ROIs, we used a group-level parcellation from a previously published data set (Fedorenko et al., 2013). We used this parcellation to generate masks for five ROIs of interest: the intraparietal sulcus (IPS), the superior precentral sulcus (sPCS), the inferior precentral sulcus (iPCS), the anterior insula/frontal operculum (AI/FO) and the inferior frontal sulcus (IFS). Because we had already defined two posterior subregions of the IPS, IPS0-1 and IPS2-3 (see *Retinotopic Mapping*), we removed all voxels belonging to these retinotopic regions from the larger IPS mask, and used the remaining voxels to define a region that we refer to as the superior IPS (sIPS). This was done within each subject separately. We intersected each subject’s mask of load-selective voxels with the mask for each ROI to generate the final MD ROI definitions.

For one subject, this procedure failed to yield any voxels in the sPCS ROI. In all analyses of sPCS, we used the remaining 9 subjects. When performing group-level ANOVA analyses of decoding performance, we performed linear interpolation based on sPCS in the remaining 9 subjects to generate an estimate of the missing value. We did this by calculating a t-score comparing the missing subject’s d’ score to the other 9 subjects in each ROI and condition where it was defined, and using the mean of these t-scores to estimate d’ in sPCS for each condition.

### Lateral Occipital Complex Localizer

We identified two subregions of the lateral occipital complex, LO and pFus, using a functional localizer developed by the Stanford Vision and Perception Lab (Stigliani, Weiner, & Grill-Spector, 2015) to identify voxels that showed enhanced response for intact objects (cars and guitars) versus phase-scrambled versions of the same images. Between 2–4 runs of this task were performed during functional scanning sessions. During each run, subjects viewed blocks of sequentially presented images in a particular category (cars, guitars, faces, houses, body parts, scrambled objects), and performed a one-back repeat detection task (mean d’ = 3.09 ± 0.49). We used a GLM to define voxels that showed significantly higher BOLD during car/guitar blocks versus scrambled blocks (FDR corrected q=0.05). We then projected this mask onto a computationally-inflated mesh of the gray matter-white matter boundary in each subject, and defined LO and pFus on this mesh based on the mask in conjunction with anatomical landmarks (Vinberg & Grill-Spector, 2008).

### Object Localizer Task

After defining the ROIs described above, voxels within each ROI were thresholded based on their visual responsiveness during performance of an independent novel object matching task. Subjects performed 2–3 runs of this task during each scanning session. This task was identical to the Identity Task described above, except that the alternate object set was used (e.g, if the subject viewed Set A during the main one-back task runs, they viewed Set B during the localizer). The object exemplars shown during this task were randomly selected for each run. Performance on this task was consistently lower than performance on the main one-back tasks, due to the fact that subjects had not been trained on this stimulus set (mean d’ = −0.23 ± 0.14).

For each subject, we combined data from all object localizer runs to generate a statistical parametric map of voxel responsiveness, based on a GLM in which all image presentations were modeled as a single predictor. We then selected only the voxels whose BOLD signal was significantly modulated by image presentation events (FDR corrected q=0.05). This limited our voxel population to those that were responsive to object stimuli that were visually similar to those presented during the main task. For one ROI in the MD network (AI-FO), this thresholding procedure yielded fewer than 10 voxels for several subjects, so for this ROI we chose to analyze the entirety of the ROI as defined by the MD localizer. The final definitions of each ROI (centers and sizes), following this thresholding procedure are summarized in Supplementary Table 1.

### MVPA Decoding

The goal of our MVPA analysis was to estimate the amount of linearly-decodable information about viewpoint match status and identity match status that was represented within each ROI during each task, and determine how this information content was affected by the task-relevance of each match dimension, as well as how it differed between correct and incorrect trials.

Before performing MVPA, we first mean-centered each single-trial voxel activation pattern, by calculating the mean across voxels on each trial, and subtracting this value from the voxel activation pattern measured on the same trial. The purpose of this was to ensure that we were measuring information related to the relative pattern of activity across voxels in each ROI, rather than information solely due to mean signal changes across conditions. Several ROIs did show a significant difference in mean signal between the identity and viewpoint tasks (see Supplementary Figure 3), however, this mean-centering step ensured that such differences did not contribute to our classification accuracy.

We performed all decoding analyses using a binary classifier based on the normalized Euclidean distance. To avoid overfitting, we used a leave-one-run-out cross-validation scheme, so that each run served as the test set once. Before starting this analysis, we removed all trials that were the first in a block, because they could not be labeled as a match or non-match, and removed all trials where the subject was incorrect or did not respond. Next, we divided data in the training set into two groups based on status as a match in the dimension of interest. For each of these two groups, we then calculated a mean voxel activation pattern (e.g., averaging the response of each voxel over all trials in the group). We also calculated the pooled variance of each voxel’s response across the two groups. Then, for each trial in the test set, we calculated the normalized Euclidean distance to each of the mean patterns of the training set groups, weighting each voxel’s contribution based on its pooled variance. We then assigned each test set trial to the group with the minimum normalized Euclidean distance. Specifically, for a training set including *n* total trials, with *n_a_* trials in condition A and *n_b_* trials in condition B, and *v* voxels in each activation pattern, we can define *ā*, 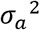, 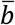, and 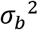 as vectors of size [1 *x v*] describing the mean and variance of each voxel’s response within conditions A and B, respectively. If *x* is a [1 *x v*] vector describing a voxel activation pattern from a single trial in the test set, the normalized Euclidean distance from *x* to each of the two training set conditions is:

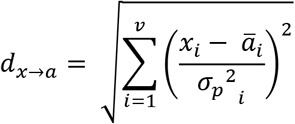

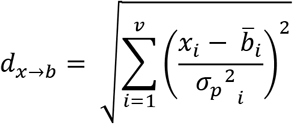

Where 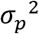 is a [1 *x v*] vector describing the pooled variance of each voxel over conditions A and B:

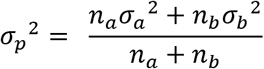

The final label assigned to each test set trial by the classifier was obtained by finding the minimum value between *d_x→a_* and *d_x→b_*. Finally, we computed a single value for classifier performance across the entire data set by calculating d’ with the formula:

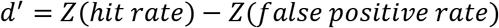

Where the hit rate is defined as the proportion of test samples in condition A accurately classified as belonging to condition A, and the false positive rate is the proportion of test samples in condition B inaccurately classified as belonging to condition A. The function *Z*(*p*), *p* ∈ [0,1] is the inverse of the cumulative distribution of the Gaussian distribution.

Because the frequency of matches in our task was less than 50%, the training set for the classifier was initially unbalanced. To correct for this, we performed downsampling on the larger training set group (non-match trials), by randomly sampling N trials without replacement from the larger set, where N is the number of samples in the smaller set. We performed 1000 iterations of this random downsampling and averaged the results for d’ over all iterations.

We assessed the significance of classifier decoding performance in each ROI using a permutation test in which we shuffled the labels of all trials in the training set and computed decoding performance on this shuffled data set. We repeated this procedure over 1000 iterations to compute a null distribution of d’ for each subject and each ROI. For each shuffling iteration, we performed downsampling to balance the training set as described above, but to reduce the computational time, we used only 100 iterations. To compute significance at the group level, we averaged the null distributions over all subjects to obtain a single distribution of 1000 d’ values, and averaged the d’ values for the real data set over all subjects to obtain a subject-average d’ value. We obtained a p-value by calculating the proportion of shuffling iterations on which the shuffled d’ exceeded the real d’, and the proportion on which the real d’ exceeded the shuffled d’, and taking the minimum value multiplied by 2. We then performed FDR correction across ROIs within each condition and match type, at the 0.01 and 0.05 significance levels (Benjamini & Yekutieli, 2001). We chose two thresholds because the 0.05 value provides slightly more power to detect weaker effects, while the 0.01 value provides a more conservative threshold.

The above analysis was carried out within each ROI, task, and match dimension separately. Next, to assess how decoding performance was affected by task, ROI, and the task-relevance of each match-dimension, we entered all d’ values into a three-way repeated measures ANOVA with factors of Task, ROI, and Relevance. Following this, to more closely investigate the interaction between ROI and Relevance, we used a nonparametric paired t-test to compare the d’ distributions for the relevant match dimension versus the irrelevant match dimension, within each task and ROI separately. This test consisted of performing 10,000 iterations in which we randomly swapped the Relevance labels corresponding to the d’ values, maintaining the subject labels. After randomly swapping, we calculated the difference in d’ between the two conditions for each subject, and used these 10 difference values to calculate a t-statistic. We then compared the distribution of these null t-statistics to the value of the t-statistic found with the real Relevance labels, and used this to generate a two-tailed p-value. These p-values were FDR corrected across ROIs.

In addition to the overall performance of the classifier, we were also interested in whether neural information about the task-relevant match dimension was associated with task performance. To evaluate this, we performed decoding of the task-relevant match dimension as described above, but here we included incorrect and no-response trials in both the training and testing sets. We considered all incorrect and no-response trials as a single group which we refer to as “incorrect”. Then, for each trial in the test set, we used the normalized Euclidean distance (calculations described above) as a metric of classifier evidence in favor of the correct trial label, where:

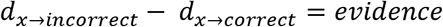

We then compared the distributions of evidence between correct and incorrect test set trials. Since there were many more correct than incorrect test set trials, we performed an additional step of downsampling to balance the training set. We divided training set trials into four groups based on their status as a match and the correctness of the subject’s response, and downsampled the number of trials in all four groups to match the number of trials in the smallest set. This was performed over 1000 iterations and the resulting values for classifier evidence were averaged. We assessed significance using a permutation test as described above. Finally, we performed a three-way ANOVA on all classifier evidence values with factors of Task, ROI, and Correctness.

In all analyses reported in the main text, we performed MVPA using all voxels from each localized ROI. Additionally, in the interest of making classification performance more comparable across ROIs, we repeated all analyses after restricting the number of voxels to 50 in each area. Overall, reducing the voxel number did not cause a dramatic change in the patterns of decoding performance across ROIs. These analyses are all reported in the Supplementary Figures 4,5, and 7.

### Image Similarity Analysis

The Viewpoint and Identity tasks were designed so that the low-level shape similarity of the images would not be explicitly informative about the status of each image as a match. Thus, when we performed classification on the status of each image as a target in each dimension, we intended to capture information that was related to perception of the abstract dimensions of each object, rather than low-level properties such as its shape in a two-dimensional projection. However, due to factors such as the small number of objects in our stimulus set, it is possible that there was some coincidental, systematic structure in the similarity between pairs of objects, such that low-level image similarity was partially informative about the status of each image as a match in either identity or viewpoint.

To evaluate this possibility, for each trial in the sequence of images shown to each subject, we determined the image similarity between the current and previous images by unwrapping each image (1000 pixels x 1000 pixels) into a single vector, and calculating the Pearson correlation coefficient between each pair of vectors. For this analysis, we removed trials that were a match in both category and viewpoint (identical images by design), and sorted the remaining trials according to whether the current and previous images were actually a match in the dimension of interest. Mean image similarity between match and nonmatch trials was compared using a one-tailed t-test (Supplementary Figure 2). This resulted in a p-value for each subject in each condition, which was used to evaluate the extent to which low-level image similarity may have been informative about match status.

In addition to using the Pearson correlation to measure similarity, we also assessed similarity by passing each image through a Gabor wavelet model meant to simulate the responses of V1 neurons to the spatial frequency and orientation content of the image (Pinto, Cox, & DiCarlo, 2008). We then compared the effectiveness of this V1 model and the simpler pixel model at capturing the responses in V1 from our fMRI data, by calculating a similarity matrix for each pairwise comparison of images (24 × 24), based on (1) the V1 model, (2) the pixel model, and (3) the voxel activation patterns recorded in V1 for each subject. We found that across all subjects, the pixel model was more correlated with the V1 voxel responses than was the V1 model (data not shown). Therefore, in the interest of parsimony, we used the simpler pixel model for all image similarity analyses.

## Acknowledgements

Funding provided by NEI R01-EY025872 to J.T.S., a James S. McDonnell Foundation Scholar Award to J.T.S. We thank Alexandra Woolgar and Kalanit Grill-Spector for code and advice regarding localizing regions of the multiple demand network and the lateral occipital complex, respectively, and Timothy Brady, Edward Vul, Vy Vo and Rosanne Rademaker for useful discussions.

## Supplementary Figures

**Supplementary Figure 1.**
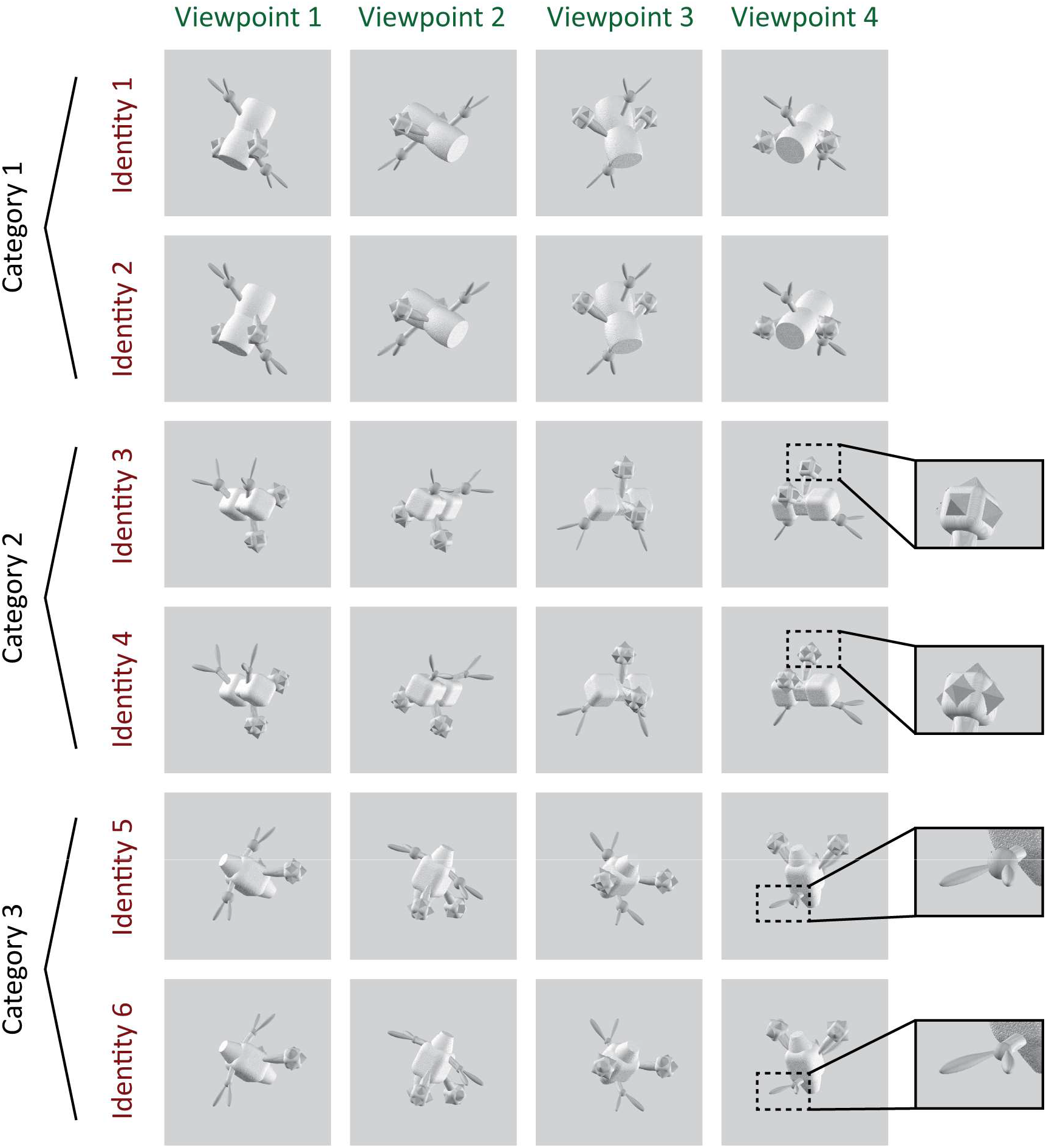
Example stimulus set from Object Set A (see Figure 1 and *Methods* for details).

**Supplementary Figure 2.**
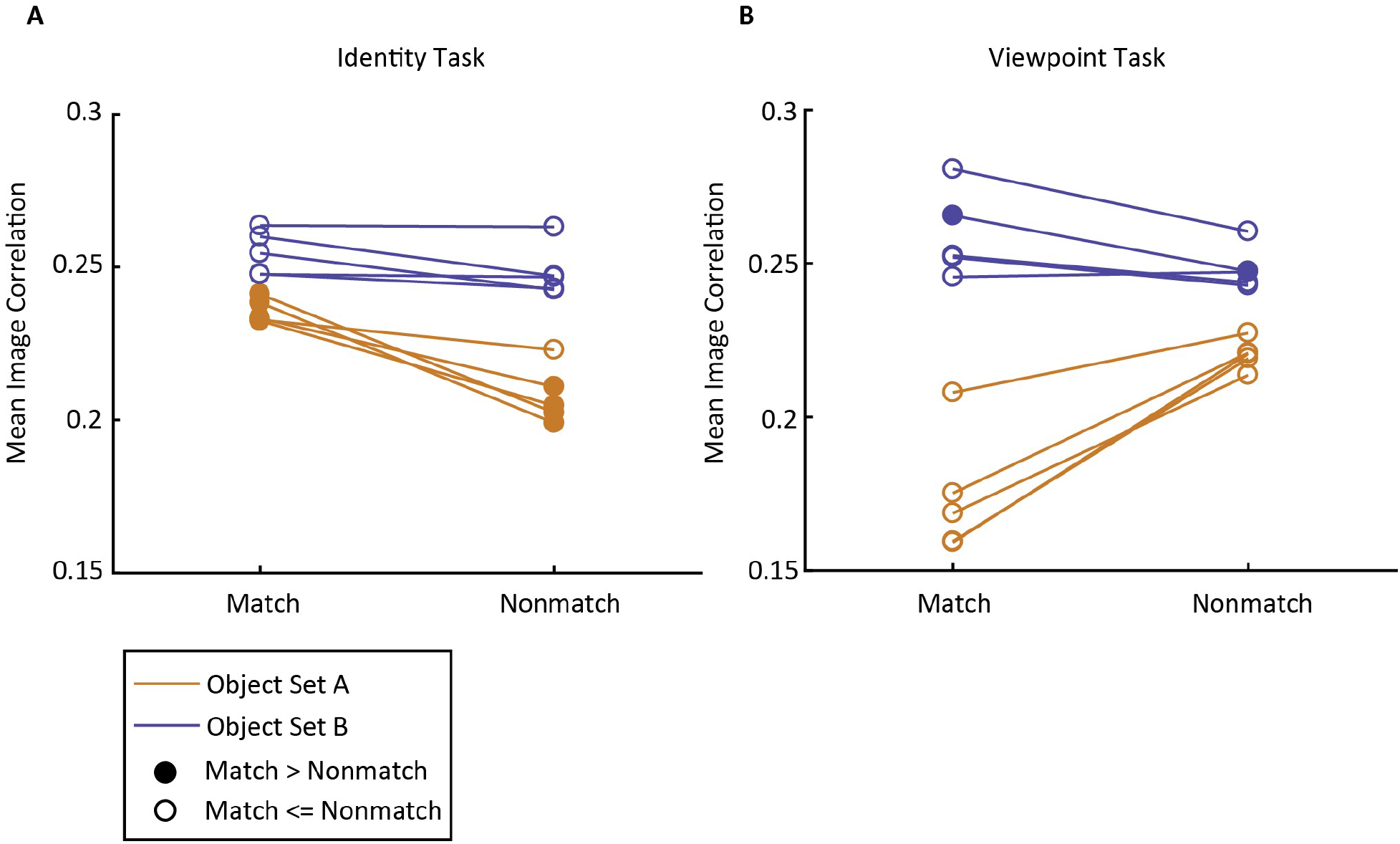
Image correlation is predictive of identity match status for several subjects in Object Set A. To assess the possibility that identity match classification in early visual cortex may have been driven by low-level similarity between pairs of images, we used the Pearson correlation coefficient to calculate the similarity between all pairs of object images (excluding image pairs that were matched in both category and viewpoint and thus highly similar). For each subject, we calculated the mean correlation coefficient between pairs of images that were a match in each feature versus images that were not a match. In the plot above, solid circles indicate that this mean value was higher for matching pairs than nonmatching pairs (α = 0.01). This finding suggests that the above-chance classification of identity match status observed in Set A subjects may have been driven by low-level image features.

**Supplementary Figure 3.**
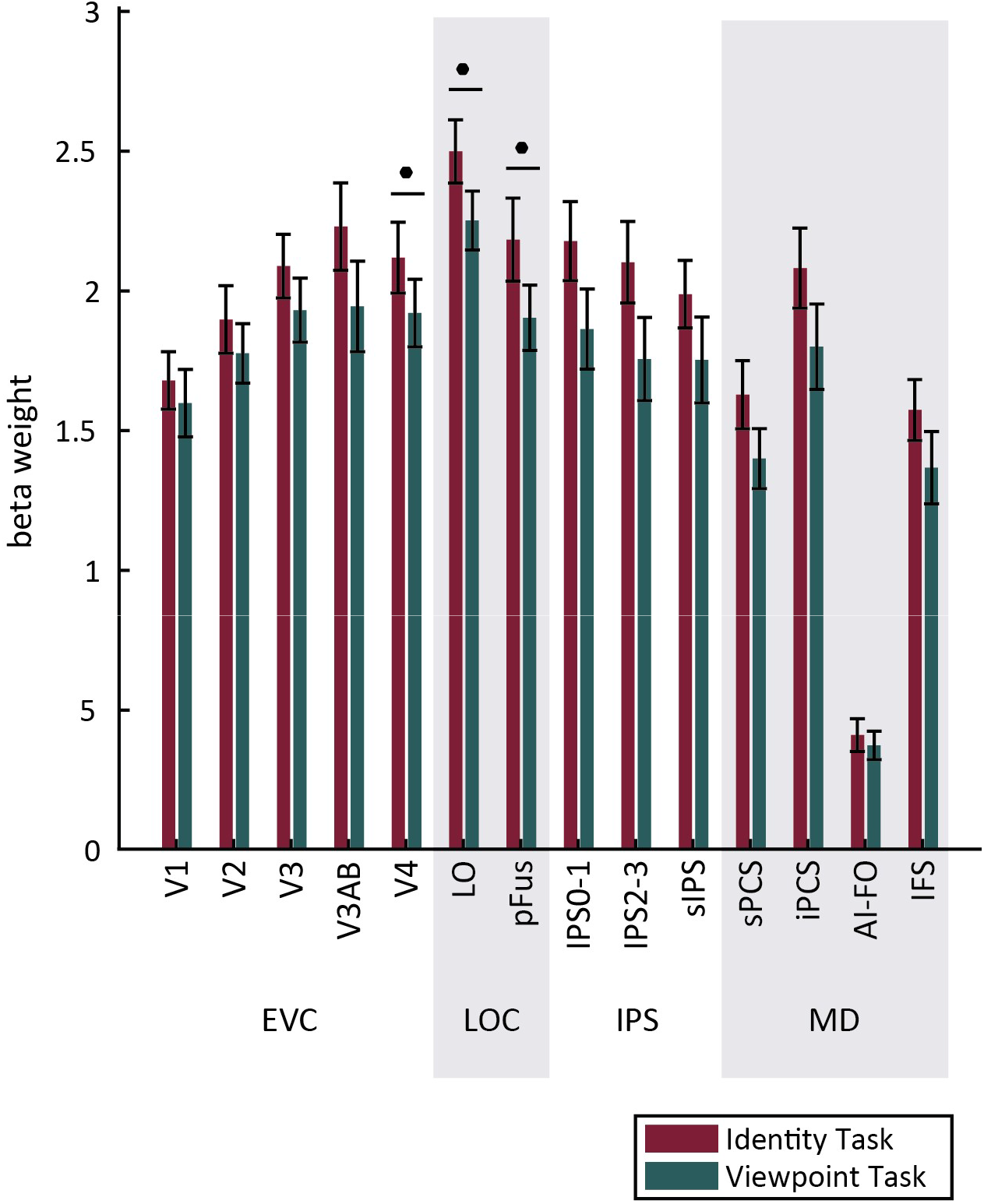
Several ROIs show a higher mean signal during the identity task than the viewpoint task. To examine whether the mean activation level in each ROI showed an effect of task, we calculated for each voxel the mean beta weight over all predictors in each condition. We then compared the mean beta weight for all voxels in each ROI between the two conditions. This revealed that for all ROIs examined, mean activation was higher during the identity task than the viewpoint task. This effect was significant in V4, LO, and pFus (bootstrapped t-test, FDR corrected, q=0.01).

**Supplementary Figure 4.**
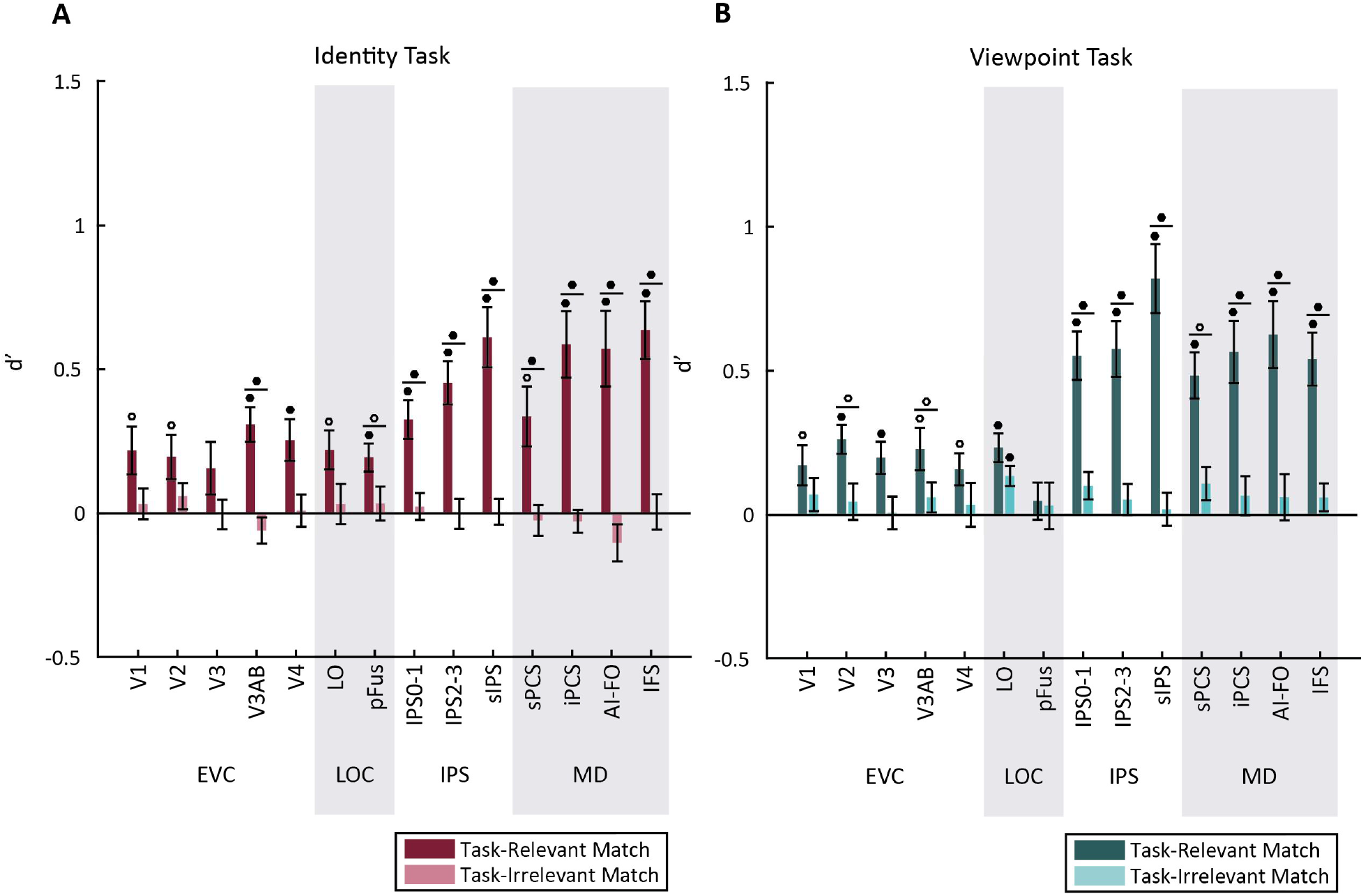
Same analysis as Figure 4, with voxel number matched across all ROIs. Voxels were selected by performing a two-sample t-test on each voxel to evaluate its selectivity for matches in the dimension of interest (e.g., the same dimension that was the target of classification), and selecting the 50 most selective voxels. To ensure that the analysis was not circular, this selection procedure was always done using only the training set data for each cross-validation fold. For one subject in area sPCS, only 32 voxels passed the localizer threshold, so for this subject and ROI we used all 32 voxels. Analysis is otherwise identical to Figure 4.

**Supplementary Figure 5.**
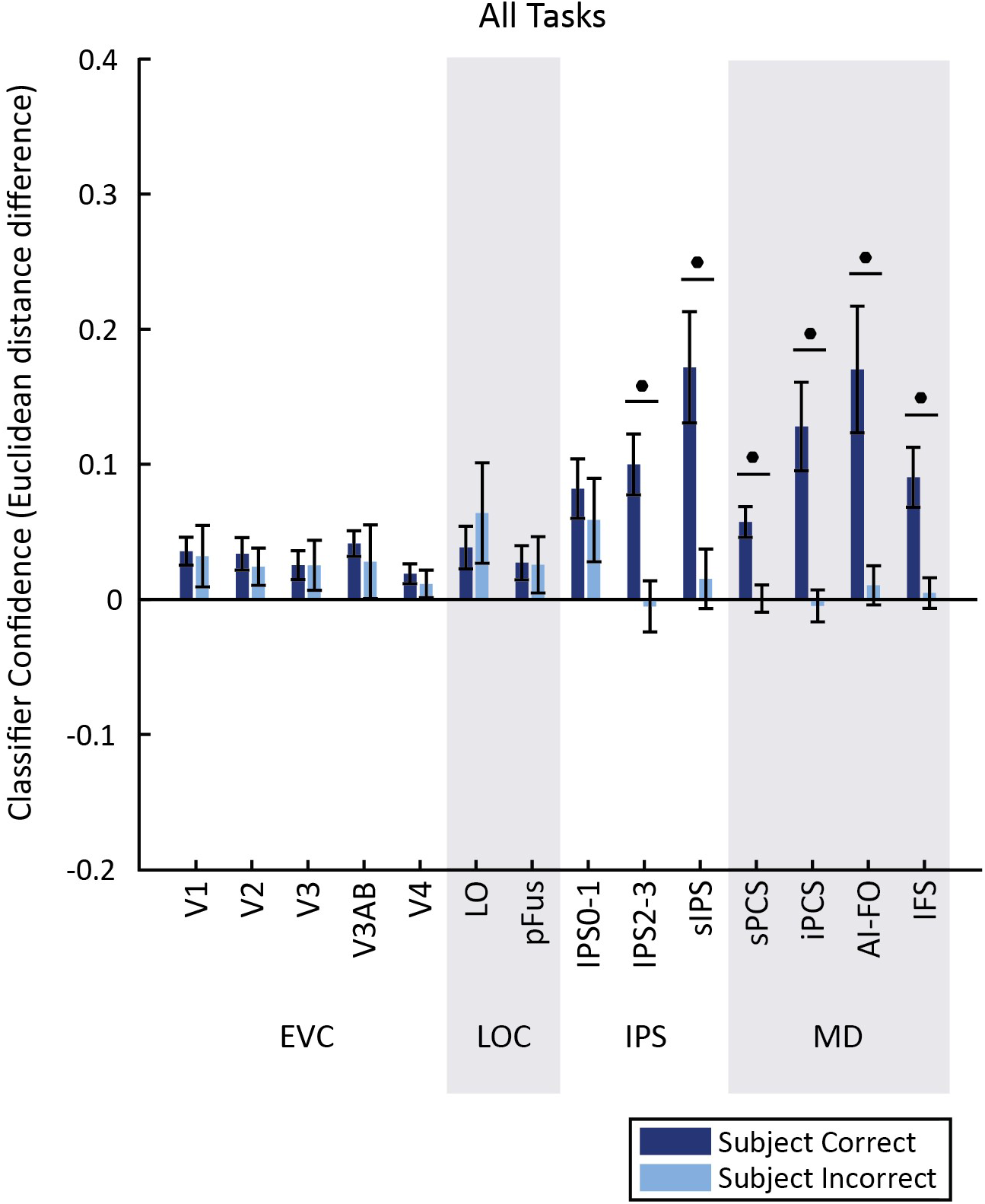
Same analysis as Figure 6, with voxel number matched across ROIs. Voxels were selected by performing a two-sample t-test on each voxel to evaluate its selectivity for matches in the dimension of interest (e.g., the same dimension that was the target of classification), and selecting the 50 most selective voxels. To ensure that there was no bias that might benefit performance on correct trials, we performed this voxel selection procedure using a training set that was balanced for correctness using downsampling over 1000 iterations, averaged the p-values for each voxel over iterations, and selected the top 50 voxels based on the resulting p-values. Analysis is otherwise identical to Figure 6.

**Supplementary Figure 6.**
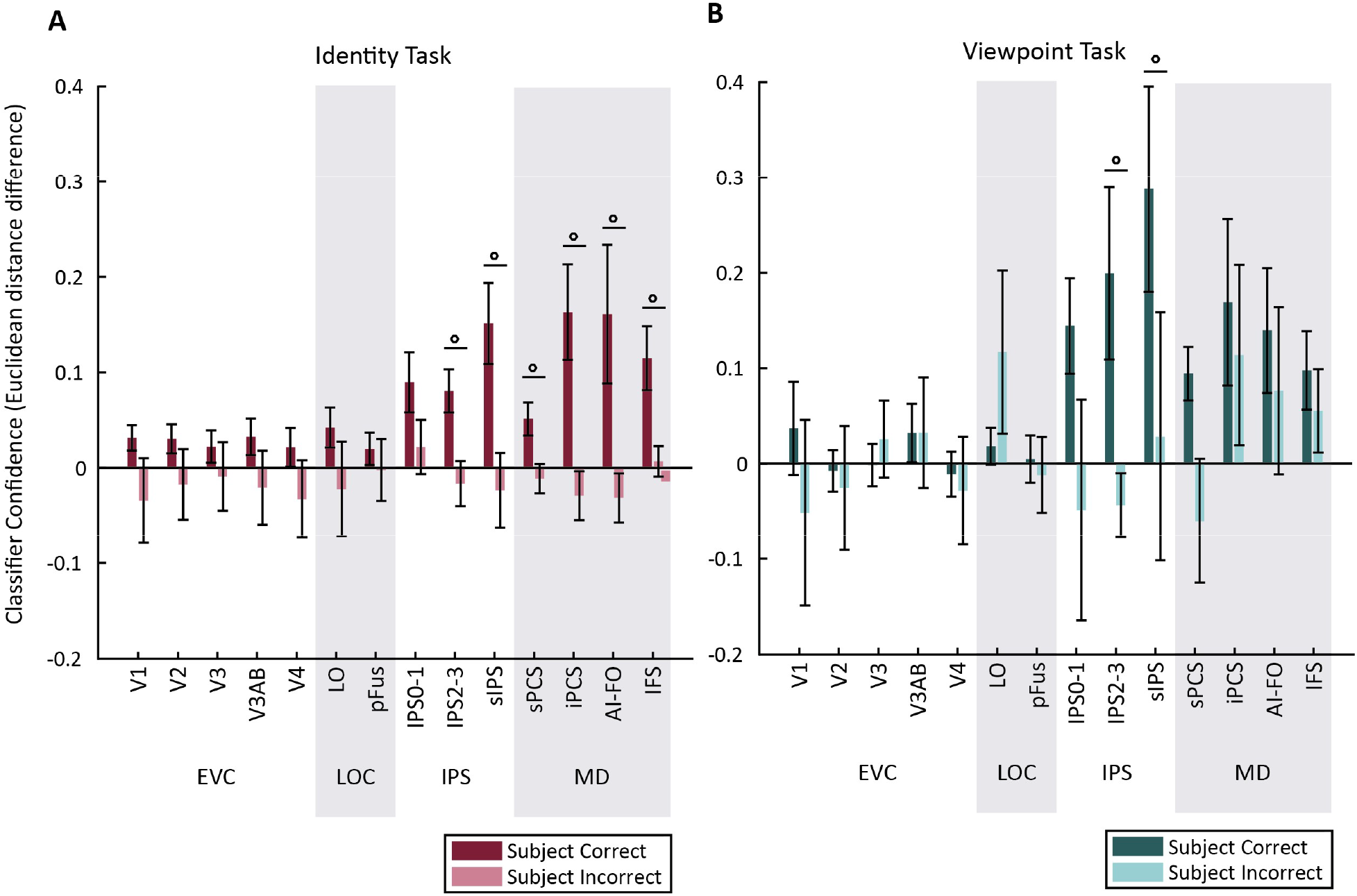
Same analysis as Figure 6, but classification was performed within each condition separately. P-values are FDR corrected across all conditions.

**Supplementary Figure 7.**
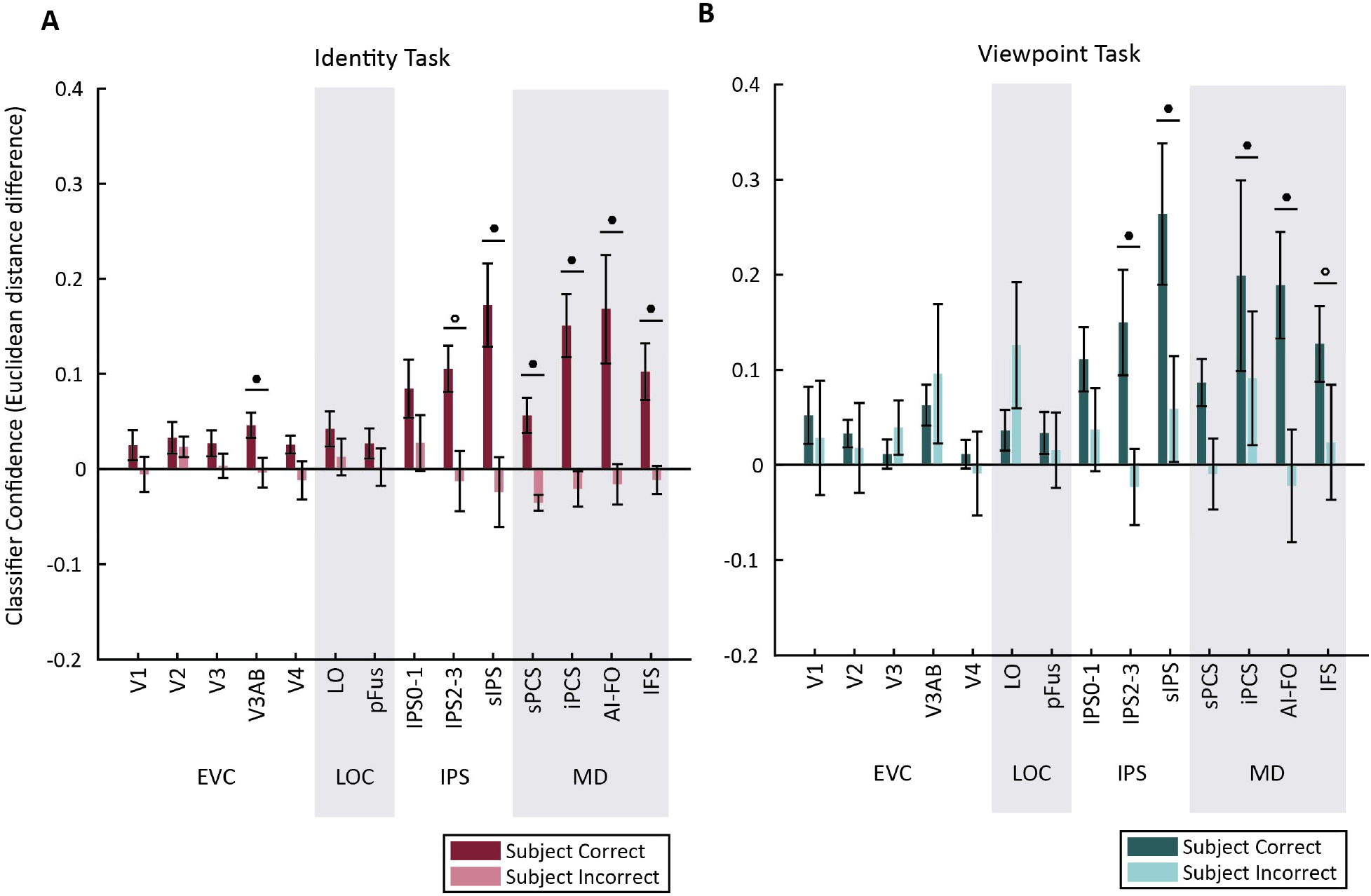
Same analysis as Supplementary Figure 6, with voxel number matched across ROIs. See Supplementary Figure 5 for details of voxel selection procedure.

**Supplementary Table 1.** Centers and sizes of the final ROIs defined for each subject, following functional localization and additional thresholding with a novel object localizer (see *Methods* for details). Coordinates of each ROI center are described in Talairach space, where X = Left-Right axis (negative is left), Y = Anterior-Posterior axis (negative is posterior), Z = Inferior-Superior axis (negative is inferior).

|  | Subject | Center |  | # Voxels |  |
| --- | --- | --- | --- | --- | --- |
|  |  | LH | RH | LH | RH |
| V1 | 1 | '[-16, -94, -12]' | '[12, -92, -17]' | 331 | 368 |
|  | 2 | '[-9, -82, 8]' | '[6, -82, 6]' | 797 | 444 |
|  | 3 | '[-15, -97, 7]' | '[11, -97, 5]' | 84 | 192 |
|  | 4 | '[-12, -97, -8]' | '[14, -94, 4]' | 249 | 186 |
|  | 5 | '[-6, -95, -4]' | '[16, -95, 1]' | 273 | 164 |
|  | 6 | '[-8, -90, -12]' | '[13, -89, -1]' | 269 | 434 |
|  | 7 | '[-9, -88, -5]' | '[11, -92, -5]' | 406 | 393 |
|  | 8 | '[-11, -92, -8]' | '[10, -91, -4]' | 348 | 254 |
|  | 9 | '[-12, -93, -8]' | '[15, -96, -5]' | 477 | 324 |
|  | 10 | '[-16, -89, -18]' | '[12, -92, -9]' | 299 | 239 |
| V2 | 1 | '[-23, -92, -11]' | '[14, -90, -15]' | 325 | 311 |
|  | 2 | '[-14, -86, 7]' | '[19, -82, 5]' | 495 | 602 |
|  | 3 | '[-17, -92, 4]' | '[15, -93, 3]' | 271 | 226 |
|  | 4 | '[-13, -93, -7]' | '[18, -92, 4]' | 354 | 212 |
|  | 5 | '[-10, -94, -8]' | '[19, -93, 5]' | 179 | 167 |
|  | 6 | '[-13, -85, -13]' | '[18, -89, -1]' | 272 | 388 |
|  | 7 | '[-14, -93, -6]' | '[15, -93, -5]' | 363 | 419 |
|  | 8 | '[-14, -87, -12]' | '[16, -89, -8]' | 416 | 308 |
|  | 9 | '[-13, -92, -6]' | '[14, -90, -6]' | 277 | 300 |
|  | 10 | '[-20, -88, -20]' | '[17, -90, -7]' | 370 | 364 |
| V3 | 1 | '[-25, -85, -8]' | '[22, -88, -9]' | 331 | 360 |
|  | 2 | '[-22, -83, 1]' | '[12, -88, 6]' | 500 | 515 |
|  | 3 | '[-19, -87, 6]' | '[21, -89, 3]' | 247 | 320 |
|  | 4 | '[-20, -90, -3]' | '[22, -85, 6]' | 359 | 216 |
|  | 5 | '[-20, -88, -7]' | '[29, -85, 1]' | 382 | 292 |
|  | 6 | '[-23, -85, -9]' | '[27, -84, -0]' | 259 | 474 |
|  | 7 | '[-24, -90, -3]' | '[23, -88, -5]' | 646 | 703 |
|  | 8 | '[-22, -83, -16]' | '[25, -88, -7]' | 432 | 348 |
|  | 9 | '[-18, -89, -3]' | '[22, -86, -6]' | 442 | 318 |
|  | 10 | '[-23, -87, -16]' | '[24, -84, -8]' | 366 | 630 |
| V3AB | 1 | '[-26, -84, 7]' | '[28, -83, 7]' | 136 | 200 |
|  | 2 | '[-28, -85, 19]' | '[23, -80, 30]' | 425 | 253 |
|  | 3 | '[-25, -84, 21]' | '[28, -85, 22]' | 269 | 214 |
|  | 4 | '[-26, -83, 11]' | '[26, -77, 19]' | 141 | 167 |
|  | 5 | '[-26, -87, 17]' | '[34, -79, 19]' | 281 | 396 |
|  | 6 | '[-31, -86, 12]' | '[31, -78, 20]' | 120 | 137 |
|  | 7 | '[-29, -83, 12]' | '[24, -78, 15]' | 357 | 306 |
|  | 8 | '[-36, -84, 5]' | '[36, -83, 1]' | 350 | 474 |
|  | 9 | '[-24, -89, 18]' | '[30, -83, 10]' | 341 | 329 |
|  | 10 | '[-25, -91, -2]' | '[28, -78, 18]' | 450 | 311 |
|  |  | LH | RH | LH | RH |
| V4 | 1 | '[-27, -75, -16]' | '[26, -75, -14]' | 116 | 245 |
|  | 2 | '[-29, -76, -8]' | '[25, -76, -8]' | 336 | 299 |
|  | 3 | '[-29, -78, -7]' | '[22, -78, -9]' | 303 | 154 |
|  | 4 | '[-22, -80, -13]' | '[25, -75, -9]' | 75 | 158 |
|  | 5 | '[-27, -79, -16]' | '[32, -75, -12]' | 243 | 139 |
|  | 6 | '[-29, -75, -17]' | '[32, -72, -14]' | 115 | 272 |
|  | 7 | '[-29, -76, -14]' | '[29, -76, -14]' | 406 | 380 |
|  | 8 | '[-31, -75, -23]' | '[30, -83, -18]' | 261 | 164 |
|  | 9 | '[-24, -76, -18]' | '[26, -77, -16]' | 304 | 362 |
|  | 10 | '[-32, -76, -22]' | '[24, -75, -17]' | 243 | 329 |
| LO | 1 | '[-43, -76, -13]' | '[37, -78, -10]' | 617 | 798 |
|  | 2 | '[-40, -77, 3]' | '[37, -78, 5]' | 911 | 1113 |
|  | 3 | '[-38, -81, 5]' | '[35, -81, 1]' | 253 | 314 |
|  | 4 | '[-33, -87, 1]' | '[34, -80, 0]' | 163 | 177 |
|  | 5 | '[-35, -84, 4]' | '[38, -81, 5]' | 207 | 132 |
|  | 6 | '[-40, -78, -7]' | '[40, -73, -3]' | 933 | 724 |
|  | 7 | '[-35, -88, -4]' | '[32, -82, -1]' | 372 | 449 |
|  | 8 | '[-41, -73, -13]' | '[38, -76, -14]' | 812 | 347 |
|  | 9 | '[-34, -80, -5]' | '[37, -78, -7]' | 792 | 634 |
|  | 10 | '[-47, -76, -14]' | '[35, -81, 2]' | 119 | 77 |
| pFus | 1 | '[-34, -75, -16]' | '[34, -51, -15]' | 215 | 54 |
|  | 2 | '[-37, -63, -10]' | '[36, -62, -9]' | 354 | 415 |
|  | 3 | '[-35, -69, -8]' | '[34, -70, -9]' | 233 | 98 |
|  | 4 | '[-37, -80, -10]' | '[36, -67, -8]' | 252 | 341 |
|  | 5 | '[-43, -66, -8]' | '[40, -64, -10]' | 201 | 127 |
|  | 6 | '[-37, -63, -16]' | '[33, -55, -15]' | 234 | 268 |
|  | 7 | '[-38, -68, -15]' | '[34, -61, -17]' | 87 | 184 |
|  | 8 | '[-37, -54, -21]' | '[32, -62, -20]' | 391 | 656 |
|  | 9 | '[-33, -63, -12]' | '[35, -61, -15]' | 96 | 447 |
|  | 10 | '[-45, -67, -12]' | '[39, -61, -15]' | 207 | 117 |
| IPS0-1 | 1 | '[-26, -80, 15]' | '[25, -79, 21]' | 162 | 236 |
|  | 2 | '[-31, -78, 27]' | '[22, -61, 40]' | 369 | 960 |
|  | 3 | '[-22, -70, 32]' | '[24, -75, 33]' | 331 | 280 |
|  | 4 | '[-22, -70, 28]' | '[25, -67, 33]' | 185 | 175 |
|  | 5 | '[-25, -70, 26]' | '[30, -67, 30]' | 524 | 397 |
|  | 6 | '[-30, -67, 26]' | '[22, -64, 40]' | 30 | 269 |
|  | 7 | '[-25, -76, 25]' | '[24, -73, 28]' | 478 | 586 |
|  | 8 | '[-34, -74, 15]' | '[32, -71, 13]' | 401 | 375 |
|  | 9 | '[-24, -75, 24]' | '[30, -74, 16]' | 675 | 482 |
|  | 10 | '[-27, -88, 9]' | '[26, -67, 33]' | 473 | 499 |
|  |  | LH | RH | LH | RH |
| IPS2-3 | 1 | '[-29, -66, 32]' | '[25, -72, 39]' | 140 | 126 |
|  | 2 | '[-25, -63, 35]' | '[24, -46, 51]' | 488 | 491 |
|  | 3 | '[-25, -54, 46]' | '[22, -61, 44]' | 283 | 203 |
|  | 4 | '[-24, -62, 45]' | '[24, -60, 39]' | 135 | 47 |
|  | 5 | '[-27, -63, 43]' | '[22, -63, 43]' | 368 | 270 |
|  | 6 | '[-26, -62, 35]' | '[24, -54, 45]' | 221 | 175 |
|  | 7 | '[-22, -71, 39]' | '[24, -61, 37]' | 395 | 352 |
|  | 8 | '[-26, -68, 27]' | '[27, -63, 23]' | 214 | 161 |
|  | 9 | '[-29, -62, 31]' | '[25, -63, 29]' | 283 | 388 |
|  | 10 | '[-27, -71, 23]' | '[25, -55, 45]' | 390 | 510 |
| sIPS | 1 | '[-34, -55, 36]' | '[34, -54, 37]' | 65 | 204 |
|  | 2 | '[-34, -40, 42]' | '[31, -47, 50]' | 155 | 283 |
|  | 3 | '[-31, -50, 43]' | '[26, -60, 45]' | 203 | 145 |
|  | 4 | '[-24, -62, 43]' | '[29, -55, 44]' | 190 | 559 |
|  | 5 | '[-32, -53, 47]' | '[34, -50, 45]' | 472 | 708 |
|  | 6 | '[-35, -48, 43]' | '[27, -51, 48]' | 527 | 434 |
|  | 7 | '[-35, -54, 41]' | '[33, -52, 44]' | 335 | 919 |
|  | 8 | '[-28, -60, 45]' | '[28, -59, 41]' | 540 | 654 |
|  | 9 | '[-34, -53, 42]' | '[32, -54, 40]' | 486 | 805 |
|  | 10 | '[-28, -55, 41]' | '[31, -51, 44]' | 1057 | 665 |
| sPCS | 1 | / | / | / | / |
|  | 2 | '[-22, 4, 60]' | '[30, 2, 54]' | 4 | 28 |
|  | 3 | '[-25, -4, 54]' | '[29, 0, 53]' | 35 | 19 |
|  | 4 | '[-31, -4, 50]' | '[36, -3, 49]' | 5 | 52 |
|  | 5 | '[-29, -4, 57]' | '[29, -4, 52]' | 56 | 142 |
|  | 6 | '[-33, 2, 54]' | '[36, 2, 55]' | 152 | 172 |
|  | 7 | '[-28, -5, 54]' | '[29, -1, 55]' | 19 | 348 |
|  | 8 | '[-37, -1, 56]' | '[29, 1, 53]' | 19 | 349 |
|  | 9 | '[-27, -3, 52]' | '[27, -3, 50]' | 189 | 209 |
|  | 10 | '[-25, -3, 56]' | '[26, 2, 55]' | 95 | 109 |
| iPCS | 1 | '[-47, 10, 34]' | '[47, 6, 35]' | 169 | 222 |
|  | 2 | '[-45, 7, 35]' | '[46, 8, 37]' | 25 | 44 |
|  | 3 | '[-48, 8, 23]' | '[47, 8, 29]' | 103 | 121 |
|  | 4 | '[-34, -2, 32]' | '[42, 6, 35]' | 30 | 217 |
|  | 5 | '[-46, 2, 32]' | '[44, 4, 28]' | 66 | 135 |
|  | 6 | '[-47, 6, 29]' | '[46, 7, 28]' | 216 | 150 |
|  | 7 | '[-42, -1, 33]' | '[41, 6, 32]' | 150 | 422 |
|  | 8 | '[-46, 8, 30]' | '[50, 5, 32]' | 124 | 154 |
|  | 9 | '[-43, 5, 33]' | '[42, 5, 34]' | 472 | 483 |
|  | 10 | '[-45, 1, 30]' | '[45, 6, 33]' | 130 | 326 |
|  |  | LH | RH | LH | RH |
| AI-FO | 1 | '[-35, 19, 4]' | '[37, 17, 5]' | 244 | 532 |
|  | 2 | '[-37, 17, 3]' | '[38, 18, 1]' | 267 | 340 |
|  | 3 | '[-35, 15, 2]' | '[32, 18, 5]' | 120 | 115 |
|  | 4 | '[-37, 18, 4]' | '[37, 18, 3]' | 354 | 510 |
|  | 5 | '[-34, 20, 1]' | '[33, 19, -1]' | 47 | 34 |
|  | 6 | '[-38, 18, 2]' | '[37, 20, 2]' | 289 | 238 |
|  | 7 | '[-35, 17, 2]' | '[35, 18, 2]' | 145 | 388 |
|  | 8 | '[-36, 16, 4]' | '[35, 17, 3]' | 218 | 485 |
|  | 9 | '[-37, 19, 4]' | '[37, 17, 5]' | 486 | 551 |
|  | 10 | '[-37, 19, 3]' | '[33, 20, 5]' | 53 | 150 |
| IFS | 1 | '[-37, 35, 28]' | '[40, 30, 28]' | 9 | 46 |
|  | 2 | '[-29, 47, 17]' | '[34, 37, 20]' | 47 | 235 |
|  | 3 | '[-44, 33, 22]' | '[42, 31, 22]' | 44 | 55 |
|  | 4 | '[-42, 29, 23]' | '[38, 30, 23]' | 5 | 96 |
|  | 5 | '[-28, 49, 21]' | '[34, 41, 18]' | 53 | 105 |
|  | 6 | '[-35, 42, 26]' | '[41, 39, 26]' | 105 | 85 |
|  | 7 | '[-40, 36, 19]' | '[40, 40, 21]' | 152 | 460 |
|  | 8 | '[-40, 33, 18]' | '[41, 33, 22]' | 71 | 308 |
|  | 9 | '[-41, 34, 26]' | '[44, 30, 27]' | 242 | 189 |
|  | 10 | '[-39, 37, 21]' | '[38, 43, 24]' | 33 | 102 |

## References

Andersson, J. L. R., Skare, S., & Ashburner, J. (2003). How to correct susceptibility distortions in spinecho echo-planar images: Application to diffusion tensor imaging. Neuroimage, 20(2), 870–888. http://doi.org/10.1016/S1053-8119(03)00336-7

Anzellotti, S., Fairhall, S. L., & Caramazza, A. (2014). Decoding representations of face identity that are tolerant to rotation. Cerebral Cortex, 24(8), 1988–1995. http://doi.org/10.1093/cercor/bht046

Benjamini, Y., & Yekutieli, D. (2001). The control of the false discovery rate in multiple testing under dependency. Annals of Statistics. Institute of Mathematical Statistics. http://doi.org/10.1214/aos/1013699998

Bichot, N. P., & Schall, J. D. (1999). Effects of similarity and history on neural mechanisms of visual selection. Nature Neuroscience, 2(6), 549–554. http://doi.org/10.1038/9205

Biederman, I. (2001). Recognizing Depth-Rotated Objects: A Review of Recent Research and Theory. Spatial Vision, 13, 241–253. Retrieved from http://www.cse.psu.edu/~rtc12/CSE597E/papers/objrecBiedermanRotationBHms.pdf

Cromer, J. A., Roy, J. E., Buschman, T. J., & Miller, E. K. (2011). Comparison of primate prefrontal and premotor cortex neuronal activity during visual categorization. Journal of Cognitive Neuroscience, 23(11), 3355–65. http://doi.org/10.1162/jocn_a_00032

Cromer, J. A., Roy, J. E., & Miller, E. K. (2010). Representation of Multiple, Independent Categories in the Primate Prefrontal Cortex. http://doi.org/10.1016/j.neuron.2010.05.005

DiCarlo, J. J., & Cox, D. D. (2007). Untangling invariant object recognition. Trends in Cognitive Sciences, 11(8), 333–341. http://doi.org/10.1016/j.tics.2007.06.010

Dubois, J., de Berker, A. O., & Tsao, D. Y. (2015). Single-unit recordings in the macaque face patch system reveal limitations of fMRI MVPA. The Journal of Neuroscience : The Official Journal of the Society for Neuroscience, 35(6), 2791–802. http://doi.org/10.1523/JNEUROSCI.4037-14.2015

Duncan, J. (2001). An adaptive coding model of neural function in prefrontal cortex. Nature Reviews Neuroscience, 2(11), 820–829. http://doi.org/10.1038/35097575

Duncan, J. (2010). The multiple-demand (MD) system of the primate brain: mental programs for intelligent behaviour. Trends in Cognitive Sciences, 14(4), 172–179. http://doi.org/10.1016/j.tics.2010.01.004

Engel, T. A., & Wang, X.-J. (2011). Same or Different? A Neural Circuit Mechanism of Similarity-Based Pattern Match Decision Making. Journal of Neuroscience, 31(19), 6982–6996. http://doi.org/10.1523/JNEUROSCI.6150-10.2011

Erez, J., Cusack, R., Kendall, W., & Barense, M. D. (2016). Conjunctive Coding of Complex Object Features. Cerebral Cortex, 26(5), 2271–2282. http://doi.org/10.1093/cercor/bhv081

Erez, Y., & Duncan, J. (2015). Discrimination of Visual Categories Based on Behavioral Relevance in Widespread Regions of Frontoparietal Cortex. Journal of Neuroscience, 35(36), 12383–12393. http://doi.org/10.1523/JNEUROSCI.1134-15.2015

Fedorenko, E., Duncan, J., & Kanwisher, N. (2013). Broad domain generality in focal regions of frontal and parietal cortex. Proceedings of the National Academy of Sciences, 110(41), 16616–16621. http://doi.org/10.1073/pnas.1315235110

Fitzgerald, J. K., Swaminathan, S. K., & Freedman, D. J. (2012). Visual categorization and the parietal cortex. Frontiers in Integrative Neuroscience, 6, 18. http://doi.org/10.3389/fnint.2012.00018

Freedman, D. J., & Assad, J. A. (2006). Experience-dependent representation of visual categories in parietal cortex. Nature, 443(7107), 85–88. http://doi.org/10.1038/nature05078

Freedman, D. J., & Assad, J. A. (2016). Neuronal Mechanisms of Visual Categorization: An Abstract View on Decision Making. Annual Review of Neuroscience, 39(1), 129–147. http://doi.org/10.1146/annurev-neuro-071714-033919

Freedman, D. J., Riesenhuber, M., Poggio, T., & Miller, E. K. (2001). Categorical representation of visual stimuli in the primate prefrontal cortex. Science, 291(5502), 312–316. http://doi.org/10.1126/science.291.5502.312

Freedman, D. J., Riesenhuber, M., Poggio, T., & Miller, E. K. (2003). A comparison of primate prefrontal and inferior temporal cortices during visual categorization. The Journal of Neuroscience : The Official Journal of the Society for Neuroscience, 23(12), 5235–5246. http://doi.org/23/12/5235 [pii]

Freiwald, W. A., & Tsao, D. Y. (2010). Functional Compartmentalization and Viewpoint Generalization Within the Macaque Face-Processing System. Science, 330(6005), 845–851. http://doi.org/10.1126/science.1194908

Grill-Spector, K., Kushnir, T., Edelman, S., Avidan, G., Itzchak, Y., & Malach, R. (1999). Differential processing of objects under various viewing conditions in the human lateral occipital complex. Neuron, 24(1), 187–203. http://doi.org/10.1016/S0896-6273(00)80832-6

Harel, A., Kravitz, D. J., & Baker, C. I. (2014). Task context impacts visual object processing differentially across the cortex. Proceedings of the National Academy of Sciences, 111(10), E962–E971. http://doi.org/10.1073/pnas.1312567111

Hayden, B. Y., & Gallant, J. L. (2013). Working memory and decision processes in visual area V4. Frontiers in Neuroscience, 7(7 FEB), 18. http://doi.org/10.3389/fnins.2013.00018

Henson, R. N. A. (2003). Neuroimaging studies of priming. Progress in Neurobiology, 70, 53–81. http://doi.org/10.1016/S0301-0082(03)00086-8

Hong, H., Yamins, D. L. K., Majaj, N. J., & Dicarlo, J. J. (2016). Explicit information for category-orthogonal object properties increases along the ventral stream. Nature Neuroscience, 19(4), 613–622. http://doi.org/10.1038/nn.4247

Ito, M., Tamura, H., Fujita, I., & Tanaka, K. (1995). Size and position invariance of neuronal responses in monkey inferotemporal cortex. Journal of Neurophysiology, 73(1), 218–226. http://doi.org/10.1152/jn.1995.73.1.218

Jackson, J., Rich, A. N., Williams, M. A., & Woolgar, A. (2017). Feature-selective Attention in Frontoparietal Cortex: Multivoxel Codes Adjust to Prioritize Task-relevant Information. Journal of Cognitive Neuroscience, 29(2), 310–321. http://doi.org/10.1162/jocn_a_01039

Jenkinson, M., Beckmann, C. F., Behrens, T. E. J., Woolrich, M. W., & Smith, S. M. (2012). Fsl. Neuroimage, 62(2), 782–790. http://doi.org/10.1016/j.neuroimage.2011.09.015

Jeong, S. K., & Xu, Y. (2016). Behaviorally Relevant Abstract Object Identity Representation in the Human Parietal Cortex. Journal of Neuroscience, 36(5), 1607–1619. http://doi.org/10.1523/JNEUROSCI.1016-15.2016

Jerde, T. A., & Curtis, C. E. (2013). Maps of space in human frontoparietal cortex. Journal of Physiology Paris, 107(6), 510–516. http://doi.org/10.1016/j.jphysparis.2013.04.002

Kosai, Y., El-Shamayleh, Y., Fyall, A. M., & Pasupathy, A. (2014). The Role of Visual Area V4 in the Discrimination of Partially Occluded Shapes. Journal of Neuroscience, 34(25), 8570–8584. http://doi.org/10.1523/JNEUROSCI.1375-14.2014

Lee, S.-H., Kravitz, D. J., & Baker, C. I. (2013). Goal-dependent dissociation of visual and prefrontal cortices during working memory. Nature Neuroscience, 16(8), 997–999. http://doi.org/10.1038/nn.3452

Lueschow, A., Miller, E. K., & Desimone, R. (1994). Inferior temporal mechanisms for invariant object recognition. Cerebral Cortex, 4(5), 523–531. http://doi.org/10.1093/cercor/4.5.523

Lui, L. L., & Pasternak, T. (2011). Representation of comparison signals in cortical area MT during a delayed direction discrimination task. Journal of Neurophysiology, 106(3), 1260–1273. http://doi.org/10.1152/jn.00016.2011

Marr, D., & Nishihara, H. K. (1978). Representation and Recognition of the Spatial Organization of Three-Dimensional Shapes. Proceedings of the Royal Society B: Biological Sciences, 200(1140), 269–294. http://doi.org/10.1098/rspb.1978.0020

Meyer, T., & Rust, N. (2018). Single-exposure visual memory judgments are reflected in inferotemporal cortex. ELife, 7, e32259. http://doi.org/10.7554/eLife.32259

Miller, E. (2000). The prefontral cortex and cognitive control. Nature Reviews Neuroscience, 1(1), 59–65. http://doi.org/10.1038/35036228

Miller, E., & Cohen, J. D. (2001). An Integrative Theory of Prefrontal Cortex Function. Annual Review of Neuroscience, 24(1), 167–202. http://doi.org/10.1146/annurev.neuro.24.1.167

Miller, E., & Desimone, R. (1994). Parallel neuronal mechanisms for short-term memory. Science, 263(5146), 520–522. http://doi.org/10.1126/science.8290960

Miller, E., Li, L., & Desimone, R. (1991). A neural mechanism for working and recognition memory in inferior temporal cortex. Science, 254(5036), 1377–1379. http://doi.org/10.1126/science.1962197

Mitchell, D. J., Bell, A. H., Buckley, M. J., Mitchell, A. S., Sallet, J., &Duncan, J. (2016). A Putative Multiple-Demand System in the Macaque Brain. The Journal of Neuroscience : The Official Journal of the Society for Neuroscience, 36(33), 8574–85. http://doi.org/10.1523/JNEUROSCI.0810-16.2016

Pagan, M., Urban, L. S., Wohl, M. P., & Rust, N. C. (2013). Signals in inferotemporal and perirhinal cortex suggest an untangling of visual target information. Nature Neuroscience, 16(8), 1132–1139. http://doi.org/10.1038/nn.3433

Pinto, N., Cox, D. D., & DiCarlo, J. J. (2008). Why is real-world visual object recognition hard? PLoS Computational Biology, 4(1), 0151–0156. http://doi.org/10.1371/journal.pcbi.0040027

Raposo, D., Kaufman, M. T., & Churchland, A. K. (2014). A category-free neural population supports evolving demands during decision-making. Nature Neuroscience, 17(12), 1784–1792. http://doi.org/10.1038/nn.3865

Riesenhuber, M., & Poggio, T. (2000). Models of object recognition. Nature Neuroscience, 3(Supp), 1199–1204. http://doi.org/10.1038/81479

Roth, N., & Rust, N. C. (2018). Inferotemporal cortex reflects behaviorally-relevant target match information during invariant object search. BioRxiv, 152181. http://doi.org/10.1101/152181

Roy, J. E., Riesenhuber, M., Poggio, T., & Miller, E. K. (2010). Prefrontal Cortex Activity during Flexible Categorization. Journal of Neuroscience, 30(25), 8519–8528. http://doi.org/10.1523/JNEUROSCI.4837-09.2010

Serences, J. T., & Saproo, S. (2012). Computational advances towards linking BOLD and behavior. Neuropsychologia, 50(4), 435–446. http://doi.org/10.1016/j.neuropsychologia.2011.07.013

Sprague, T. C., & Serences, J. T. (2013). Attention modulates spatial priority maps in the human occipital, parietal and frontal cortices. Nature Neuroscience, 16(12), 1879–1887. http://doi.org/10.1038/nn.3574

Stigliani, A., Weiner, K. S., & Grill-Spector, K. (2015). Temporal Processing Capacity in High-Level Visual Cortex Is Domain Specific. Journal of Neuroscience, 35(36), 12412–12424. http://doi.org/10.1523/JNEUROSCI.4822-14.2015

Swisher, J. D., Halko, M. A., Merabet, L. B., McMains, S. A., & Somers, D. C. (2007). Visual Topography of Human Intraparietal Sulcus. Journal of Neuroscience, 27(20), 5326–5337. http://doi.org/10.1523/JNEUROSCI.0991-07.2007

Tanaka, K. (1993). Neuronal mechanisms of object recognition. Science (New York, N.Y.), 262(5134), 685–8. Retrieved from http://www.ncbi.nlm.nih.gov/pubmed/8235589

Tanaka, K. (1996). Inferotemporal cortex and object vision. Annual Review of Neuroscience, 19, 109–39. http://doi.org/10.1146/annurev.ne.19.030196.000545

Tarr, M. J., Williams, P., Hayward, W. G., & Gauthier, I. (1998). Three-dimensional object recognition is viewpoint dependent. Nature Neuroscience, 1(4), 275–277. http://doi.org/10.1038/1089

Turk-Browne, N. B., Yi, D. J., Leber, A. B., & Chun, M. M. (2007). Visual quality determines the direction of neural repetition effects. Cerebral Cortex, 17(2), 425–433. http://doi.org/10.1093/cercor/bhj159

Vaziri-Pashkam, M., & Xu, Y. (2017). Goal-Directed Visual Processing Differentially Impacts Human Ventral and Dorsal Visual Representations. The Journal of Neuroscience, 37(36), 8767–8782. http://doi.org/10.1523/JNEUROSCI.3392-16.2017

Vinberg, J., & Grill-Spector, K. (2008). Representation of Shapes, Edges, and Surfaces Across Multiple Cues in the Human Visual Cortex. Journal of Neurophysiology, 99(3), 1380–1393. http://doi.org/10.1152/jn.01223.2007

Wandell, B. A., Dumoulin, S. O., & Brewer, A. A. (2007). Visual field maps in human cortex. Neuron, 56(2), 366–383. http://doi.org/10.1016/j.neuron.2007.10.012

Waskom, M. L., Kumaran, D., Gordon, A. M., Rissman, J., & Wagner, A. D. (2014). Frontoparietal Representations of Task Context Support the Flexible Control of Goal-Directed Cognition. Journal of Neuroscience, 34(32), 10743–10755. http://doi.org/10.1523/JNEUROSCI.5282-13.2014

Winawer, J., & Witthoft, N. (2015). Human V4 and ventral occipital retinotopic maps. Visual Neuroscience, 32, E020. http://doi.org/10.1017/S0952523815000176

Woloszyn, L., & Sheinberg, D. L. (2009). Neural Dynamics in Inferior Temporal Cortex during a Visual Working Memory Task. Journal of Neuroscience, 29(17), 5494–5507.http://doi.org/10.1523/JNEUROSCI.5785-08.2009

Woolgar, A., Thompson, R., Bor, D., & Duncan, J. (2011). Multi-voxel coding of stimuli, rules, and responses in human frontoparietal cortex. Neuroimage, 56(2), 744–752. http://doi.org/10.1016/j.neuroimage.2010.04.035

